# On the ecology of *Acinetobacter baumannii* – jet stream rider and opportunist by nature

**DOI:** 10.1101/2024.01.15.572815

**Authors:** Gottfried Wilharm, Evelyn Skiebe, Andżelina Łopińska, Paul G. Higgins, Kristin Weber, Christoph Schaudinn, Christof Neugebauer, Katharina Görlitz, Gideon Meimers, Yana Rizova, Ulrike Blaschke, Christine Heider, Christiane Cuny, Stephan Drewes, Elisa Heuser, Kathrin Jeske, Jens Jacob, Rainer G. Ulrich, Marcin Bochenski, Mariusz Kasprzak, Ewa Burda, Mateusz Ciepliński, Ireneusz Kaługa, Łukasz Jankowiak, José I. Aguirre, Alejandro López-García, Ursula Höfle, Zuzanna Jagiello, Marcin Tobółka, Bartosz Janic, Piotr Zieliński, Maciej Kamiński, Johannes Frisch, Joachim Siekiera, Andreas F. Wendel, Paul Brauner, Udo Jäckel, Michael Kaatz, Stefanie Müller, Antina Lübke-Becker, Lothar H. Wieler, Johanna von Wachsmann, Lakshmipriya Thrukonda, Mustafa Helal, Lennard Epping, Silver A. Wolf, Torsten Semmler, Leszek Jerzak

**Author notes:** Correspondence to: Gottfried Wilharm, Robert Koch-Institut, Projektgruppe P2, Bereich Wernigerode, Burgstr. 37, D-38855 Wernigerode, Germany.

## Abstract

The natural reservoirs of the nosocomial pathogen *Acinetobacter baumannii* are not well defined. We previously identified white storks as a model system to study the ecology of *A. baumannii*. Having screened more than 1,300 white stork nestlings over a period of six years across different regions of Poland and Germany (overall isolation rate of ∼29.5%), including food chain analyses and environmental samplings, we come up with a detailed picture of the dynamics and diversity of *A. baumannii* in their natural habitats. Adult storks, rather than being stably colonized with strains of *A. baumannii* which are successively transferred to their offspring, instead initially encounter these bacteria while foraging. Among their common food sources, consisting of earthworms, small mammals, and insects, we identified earthworms as a potential source of *A. baumannii*, but more so the associated soil as well as plant roots. Through this, hotspot soil and compost habitats were identified which enable population dynamics to be studied over the course of the year. We demonstrate that sterilized plant material is rapidly colonized by airborne *A. baumannii* suggesting they patrol to search for novel habitats, being opportunist by nature. The prevalence of *A. baumannii* exhibited a strong seasonality and peaked during summer. The strains we collected in Poland and Germany represent more than 50% of the worldwide known diversity in terms of the intrinsic OXA-51-like β-lactamase. A set of ∼400 genomes was determined and compared to a diverse set of publicly available genomes. Our pan-genome estimate of the species (∼51,000 unique genes) more than doubles the amount proposed by previous studies. Core-genome based phylogenetic analyses illustrated numerous links between wildlife isolates and hospital strains, including ancient as well as recent intercontinental transfer. Our data further suggest massive radiation within the species early after its emergence, matching with human activity during the Neolithic. Deforestation in particular seemed to set the stage for this bloom as we found that forests do not provide conducive conditions for the proliferation of *A. baumannii*. In contrast, wet and nutrient-rich soil alongside rivers sampled during the summer can yield an isolation rate of ∼30%. Linking published work on the interaction between *A. baumannii* and fungi and on aspergillosis as a major cause of mortality in white stork nestlings to our findings, we hypothesized that fungi and *A. baumannii* share a long history of coevolution. Interaction studies revealed the capability of *A. baumannii* to adhere to fungal spores and to suppress spore germination. Taken together, the intrinsic resistance endowment and potential to acquire antibiotic resistance can be explained by coevolution with antibiotic-producing fungi and other microorganisms within soil, and resistance to desiccation stress and radiation can be interpreted in the light of intercontinental hitchhiking through fungal spores.

**Originality - Significance:** The ecology of the nosocomial pathogen *Acinetobacter baumannii* remains poorly understood outside the hospital. Here, we present the most comprehensive study on its environmental biology to date, after having collected more than 1,450 independent isolates of which around 400 were whole genome-sequenced. This study more than doubles the size of the pan-genome of the species, illustrating both the diversity of our collection and the bias of previous work, but also the bottleneck for the establishment of lineages within the hospital environment. We reached isolation rates of about 30% both in white stork (*Ciconia ciconia*) nestlings and in soil samples when considering for sampling all preferences of *A. baumannii* we uncovered. Thus, it is now possible to study the ecology and evolution of *A. baumannii* in nature at an unprecedented temporal and spatial resolution. We describe the worldwide spread of *A. baumannii* lineages in nature as an ancient phenomenon that even surpasses that of human-associated bacteria in magnitude. This is likely due to airborne spread, putatively facilitated by association with fungal spores. We propose that *A. baumannii* is an opportunist by nature, using airborne patrolling to rapidly enter new suitable habitats consisting of organic matter in early stages of decomposition. Our collective data suggest that *A. baumannii*, early after its speciation, went through massive radiation during the Neolithic, likely due to deforestation, settlement and farming producing numerous favorable habitats. Their natural lifestyle, which requires rapid adaptability to various habitats as well as tolerance to desiccation, radiation and antibiotic stress, perfectly predispose these opportunistic pathogens to establish within the hospital setting. Comparison of genomes from environmental and clinical isolates will now enable studies of the adaptive evolution of environmental bacteria towards multidrug-resistant opportunistic pathogens.

## Introduction

The Gram-negative *Acinetobacter baumannii* is notorious for its potential to act as an opportunistic pathogen in hospitals, facilitated by an exceptional resistance to desiccation stress and intrinsic as well as acquired resistance to antibiotics and disinfectants. The genus *Acinetobacter* currently comprises of around 100 described or tentative species (80, 83), the majority of which consist of environmental species, or commensals of human and animals that rarely cause any disease. Some species like *A. lwoffii* and *A. johnsonii*, which are regularly found on human skin, mucosa and within the gut of humans and animals, appear to play an essential role in the development of immune tolerance (29, 37, 98, 99). In contrast, the species most frequently involved in hospital infections, e.g. *A. baumannii*, *A. pittii*, *A. nosocomialis*, and *A. seifertii* do not represent commensals of humans and animals, and their natural habitats are still poorly described (9, 81, 102). Although the zoonotic potential of *A. baumannii* is evident from previous work (25, 56, 68, 95, 129), the pathogen is not generally accepted as zoonotic, e.g. not listed as zoonotic pathogen by public health agencies such as the Centers for Disease Control and Prevention (CDC), Public Health England (PHE) or the European Centre for Disease Prevention and Control (ECDC) (https://www.cdc.gov/media/releases/2019/s0506-zoonotic-diseases-shared.html; https://www.gov.uk/government/publications/list-of-zoonotic-diseases/list-of-zoonotic-diseases; https://www.ecdc.europa.eu/sites/default/files/documents/j-efsa-2021-6971.pdf). As a hallmark, the killing of more than 400 individuals of a flock of sheep by a multidrug-resistant (MDR) and hypervirulent strain related to international clone 2 (IC2) illustrates the threat of the bacterium shuttling between human and animal host systems (63). The relevancy to One-Health is further underlined by the reservoir of antibiotic resistance genes found in environmental *Acinetobacter* spp., as well as other bacteria, that can be mobilized into pathogenic *Acinetobacter* spp. via horizontal gene transfer (33, 41, 133). Moreover, the release of MDR *A. baumannii* from wastewater into the environment has been well-described (42, 51).

Here, starting from the association of *A. baumannii* with white stork nestlings as recently described (129), we intended to unravel the ecology of these bacteria in their natural habitats.

## Results

### Update on the prevalence of *A. baumannii* in white stork nestlings

Previously, we had presented data from 661 white stork nestlings sampled between 2013 and 2016 in Poland which were found to be colonized with *A. baumannii* in the choana region at an average rate of 25% (129). We continued the systematic study of this environment and, based on 1,319 white stork nestlings sampled between 2013 and 2018 across multiple regions of Poland, the average colonization rate was identified as 29.5% (Fig. 1; Suppl. Table S1). White stork nestlings were also sampled on a smaller scale in Germany in 2015 and the positive rate of choana samples reached 34.5% (n=29). Choana samplings in Spain near Ciudad Real in 2015 revealed no positives from 57 white stork nestlings, but a positive rate of 7.8% (n=64) was detected among white stork nestlings sampled in 2019 near Madrid (Suppl. Table S1). Interestingly, this included for the first time the isolation of *A. nosocomialis* (n=2) from white stork samples, indicating differences in the ecology of bacteria and/or storks in Spain compared to Poland and Germany.

**Fig. 1:**
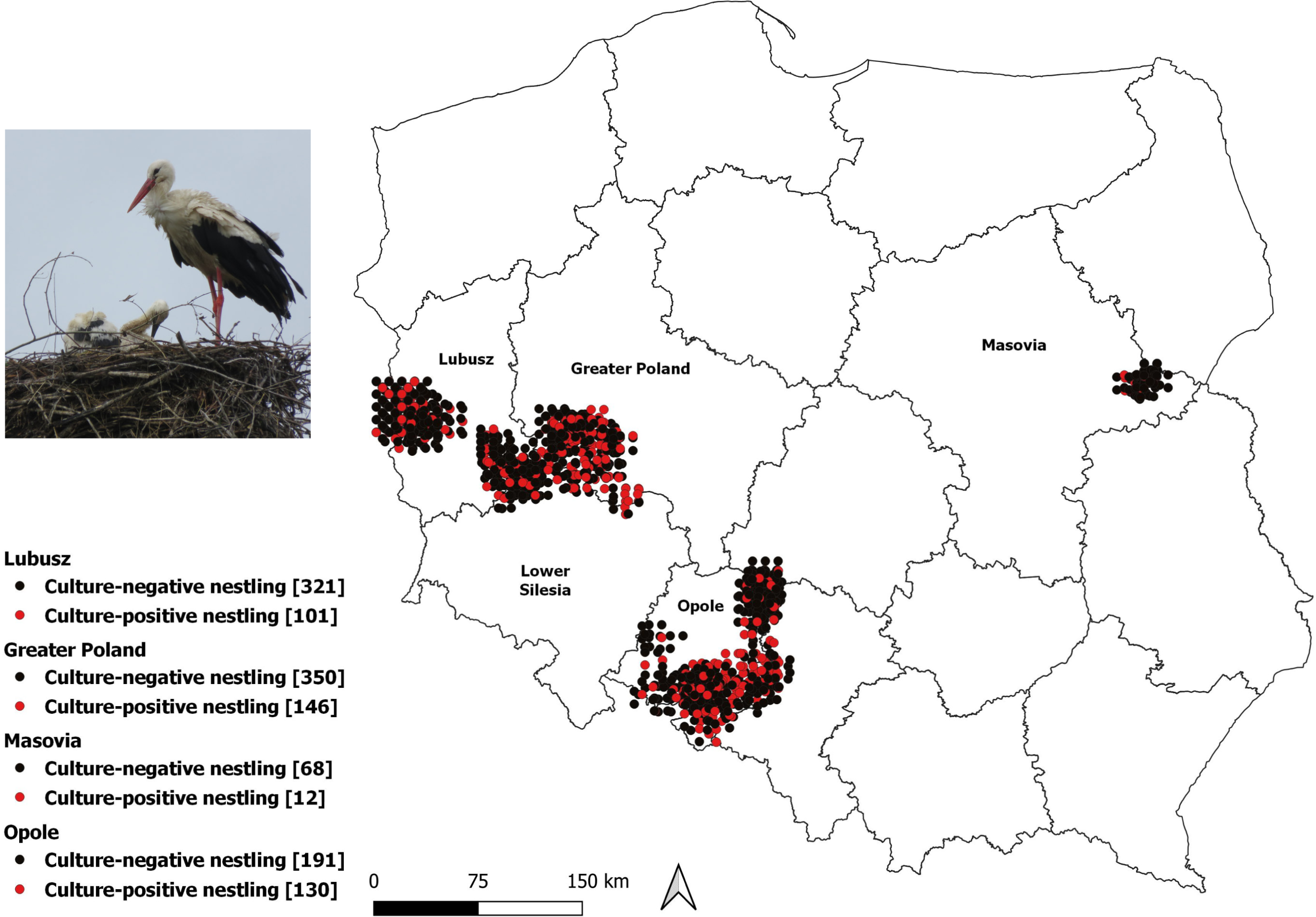
Map of the sampling sites of white stork nestlings in the years 2013-2018 in Poland. The overall positive rate for isolation of *A. baumannii* from choana swabs of nestlings was 29.5% (n=1,319). Nestlings tested positive for *A. baumannii* are represented by red circles, nestlings tested negative are indicated black.

Rectal sampling of white stork nestlings was conducted on 747 individuals in different regions of Poland throughout the study period and revealed an overall positive rate of 8%, however with great variability depending on the region and year of sampling (Suppl. Table S1). Rectal sampling of 64 white stork nestlings in Spain in 2019 yielded no *A. baumannii* isolates. Thus, choana sampling yielded higher isolation rates of *A. baumannii* compared to rectal sampling.

Notably, colonization of white stork nestlings by *A. baumannii* was apparently not associated with increased mortality of the chicks. However, significantly increased white blood cell counts in colonized nestlings argue for an infection by *A. baumannii* rather than mere colonization (Suppl. Table S2).

### Pellets from white storks

We had previously described that white stork spit pellets, consisting of indigestible remains of their food, are occasionally contaminated with *A. baumannii* and might therefore be of interest for studying the ecology of both bacteria and storks (129). We collected pellets from selected nests in Loburg and neighbouring villages (Germany/state of Saxony-Anhalt) during March to August throughout the years 2015 and 2016. Interestingly, pellet samples proved positive for *A. baumannii* not earlier than the end of May in 2015 and the end of April in 2016, respectively (Suppl. Fig. S1). In 2015, positive samples were identified until migratory departure of the storks in August, with an overall rate of 14.8% positive pellets (13 out of 88), while in 2016 only a few positive samples were found from end of April until June (overall 6.9% positive pellets, 6 out of 87). Sporadically collected egg shells and feathers were also a source of *A. baumannii* in few cases, and a stork chick found dead was also sampled positive in 2016. Pellets collected from stork colonies in Spain during the winter season of 2015 were negative irrespective of their particular feeding grounds (landfills or nature, Suppl. Fig. S2). Overall, our data suggest that white storks arriving from their wintering grounds do not appear to be stably colonized by *A. baumannii*. Rather, they seem to acquire these bacteria after arrival in the summer quarters depending on the progression of the season, likely via the food chain.

It is worth mentioning, that pH values in pellets contaminated with *A. baumannii* were found to be as low as pH 2 in some cases and it was proven that the bacteria were not only localized at the surface of the pellets but also inside (Suppl. Fig. S3). Consequently, although under laboratory conditions *A. baumannii* cannot thrive at pH 2, it can survive such harsh conditions in a natural setting.

### Diversity of strains in individual nests

To challenge our hypothesis on an only transient carriage/colonization of parental white storks with *A. baumannii* we analyzed the diversity of strains isolated from nestlings within individual nests. As a proxy of diversity, we made use of the intrinsic *bla*_OXA-51-like_ gene of *A. baumannii* encoding the class D oxacillinase OXA-51, the protein variants of which are indexed and previously proven useful as phylogenetic marker (95, 121, 129, 135). We sequenced the *bla*_OXA-51-like_ gene of all isolates collected from white stork nestlings within a single year in a selected region of Poland and visualized the diversity on a map (Fig. 2). The diversity found within individual nests was striking, with up to 5 nestlings each carrying a distinguishable strain. These findings are inconsistent with a vertical transmission from stably colonized parental storks and support our hypothesis of a transient colonization/carriage via food intake from a diverse *A. baumannii* population.

**Fig. 2:**
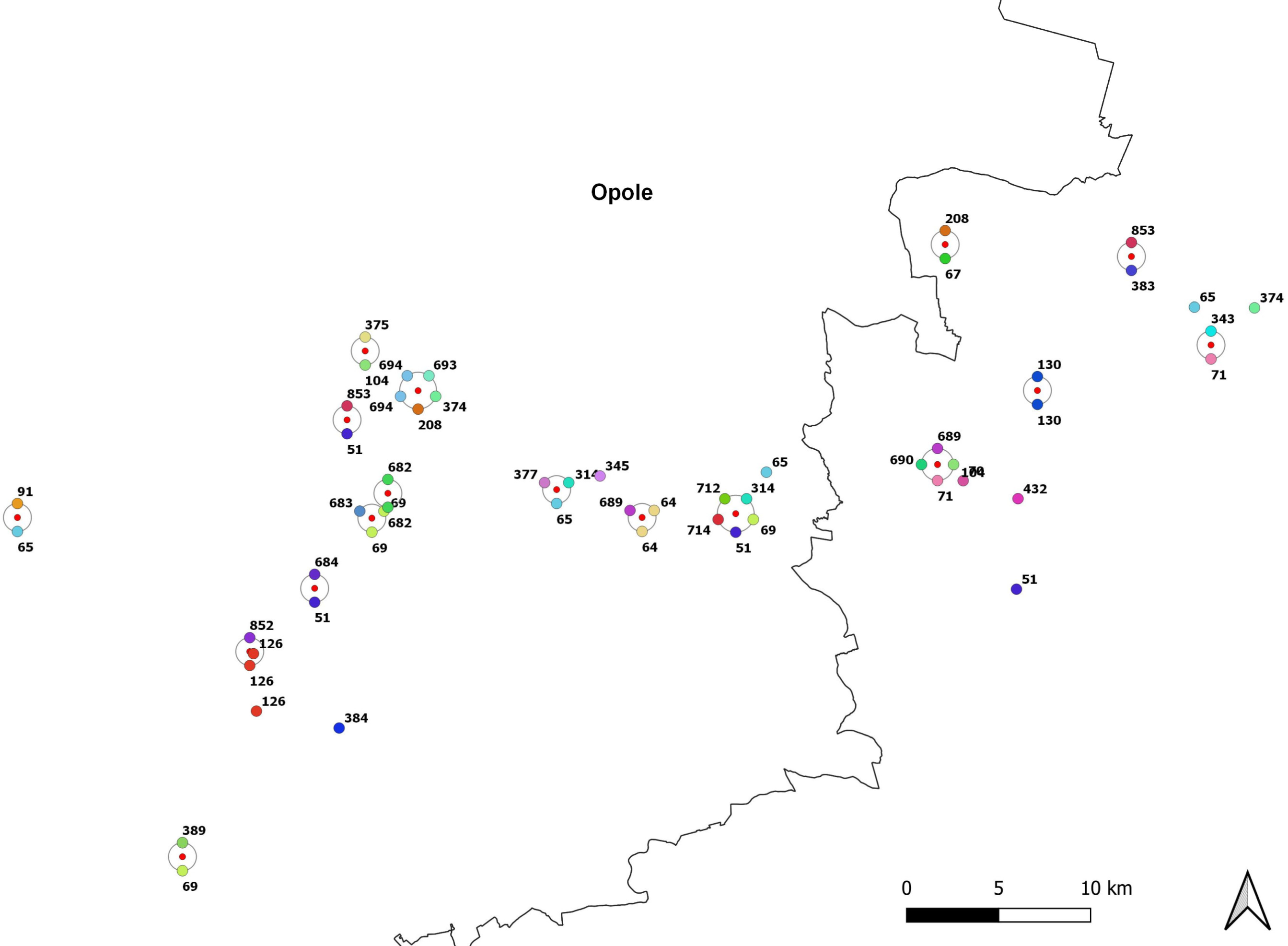
Diversity of *A. baumannii* isolates within individual nests. Cartographic presentation of individual nests with nestlings tested positive for *A. baumannii* within a region of Opole in the year 2016. Nests with more than one nestling tested positive are represented by a central red-filled circle. The numbers represent OXA-51 variant numbers. Nestlings tested negative have not been visualized for the sake of clarity.

### Tracing the food sources

The predominant prey of white storks consists of small vertebrates, especially rodents and shrews, as well as arthropods and earthworms (11, 58). In line with the literature, the dominant contribution of rodents and arthropods to the diet of white storks could be directly estimated from pellets of white storks collected in Poland and Germany (Suppl. Fig. 4), although analysis of the remains of earthworms from pellets, while possible (87), is time-consuming. To assess the potential role of beetles and other insects as sources of *A. baumannii*, we compared the content of pellets tested positive for *A. baumannii* with that of pellets tested negative and found no difference in the fraction of insect matter (Suppl. Fig. S5). Moreover, dominant arthropods found in pellets such as the ground beetles *Zabrus tenebrioides* and *Pterostichus melanarius* (Suppl. Fig. 6) collected from nature, albeit at low numbers (n=13), were also found negative for *A. baumannii*. We also had the opportunity to sample grasshoppers collected in great quantities by an individual, deceased white stork, which was dissected and found gorged with grasshoppers, however all of the swallowed grasshoppers (>50) tested negative for *A. baumannii*. Moreover, pellets from Spanish storks, containing high amounts of crab and grasshopper remains (Suppl. Fig. S2), also tested negative. Altogether, we found no evidence of a contribution of arthropods captured by white storks to the acquisition of *A. baumannii*.

Next, we examined the potential role of rodents and shrews as a source of *A. baumannii*. We collected cat-captured rodents and shrews throughout Germany (states of Saxony-Anhalt, Hesse and Thuringia) in the period from 2014 to 2017 and took samples from the trachea and rectum. We found an average colonization rate of 2% for rodents (n=154) and 4.7% for shrews (n=64). Interestingly, all individuals colonized with *A. baumannii* were captured between August and September (Suppl. Fig. S7). Next, we sampled wild, captive and laboratory rats (*Rattus norvegicus*) from Germany (75, 94, 109), as well as other rodents of the genera *Microtus*, *Myodes*, and *Apodemus* captured in Germany (48, 49) in the period 2007 to 2017. No rats and other rodents were found positive (n_rats_=491 and n_other rodents_=532).

Earthworms were collected in Germany in the state of Saxony-Anhalt between August 2016 and September 2020. Out of 618 individual earthworms in total, we isolated *A. baumannii* from 22 earthworms (3.6%) provisionally assigned to the genera *Lumbricus*, *Octolasium*, *Dendrobaena* and *Eisenia* (Suppl. Tab. 1). Of these positive samples, 19 were collected from May to September and only three isolates dated from March, April and November, respectively (1% positive of n=75 earthworms collected between October 21^st^ and March 21^st^). The overall positive rate of *A. baumannii* isolated from garden compost earthworms collected in Wernigerode (Germany) was 4.2% (14/331), and 4.3% (4/94) for a specific site at the bank of the river Holtemme (Germany), where we had recently also isolated *A. nosocomialis* from soil (128).

### Other birds and their prey

We reasoned that studying other birds with an overlapping prey spectrum compared to white storks could help to indicate specificities of the relationship to *A. baumannii*. Consequently, we collected pellets, feathers and egg shells from a breeding colony of grey heron (*Ardea cinerea*) in Germany. With the exception of one egg shell sample collected in June from which we could isolate *A. baumannii* (Suppl. Fig. S8), all samples (n=35) were negative. The detailed analysis of the pellets indicated their diet to consist of a significant portion of small mammals and insects (Suppl. Fig. S8). Further, we collected pellets (n=101) from kestrel (*Falco tinnunculus*) between March and October 2015 and 2016, and found three isolates (3%) of *A. baumannii*, each collected between August and September (Suppl. Fig. S9). Occasionally discovered owls’ pellets (n=6), putatively from *Tyto alba* and *Asio otus*, collected in August 2015, were also positive for *A. baumannii* in two cases. Moreover, we took rectal samples from nestlings of the black stork (*Ciconia nigra*), a species which, in contrast to white stork, is not synanthropic and is preferentially breeding and foraging in forests. Notwithstanding, anthropogenic waste was found in 26% of occupied nests (45). The diet of black stork nestlings in Poland is known to be dominated by fish and amphibians (53). Not a single one of 64 rectal samples collected in Poland from black stork nestlings over a period of four years was found positive for *A. baumannii*. Taken together, grey heron, kestrel and black stork differ significantly from white stork regarding carriage with *A. baumannii*.

### Compost, soil, rhizosphere and the forest paradox

Following up on a possible role of earthworms in the transmission of *A. baumannii* to white storks, we started collecting soil samples. Initially, we selected compost soil which is wet and rich in decomposing material, providing an environment from which earthworms could be easily collected. As we realized early on that compost soil represented an excellent source for *A. baumannii*, we utilized a boring rod to study the profile within one meter depth of the compost (Suppl. Fig. S10). Continued sampling of the soil originating from a single compost across a period of seven months yielded 86 isolates containing no less than 20 different variants of the OXA-51 family (60/91 samples positive, 66%). Concomitant sampling of earthworms within the same compost yielded 10 different variants of OXA-51 from 7/44 (16%) positive earthworms. We observed a remarkable dynamic of isolated lineages and of the depth-depending colonization over time. The compost, which was continuously used for the deposition of vegetable waste and egg shells during this period, was not permanently colonized throughout all layers and there was a shift of the dominant lineages isolated over time. Continued sampling in subsequent years revealed that isolation rates declined (i) during winter and (ii) within 8-10 weeks after stopping to replenish the compost with fresh material.

Due to the anthropogenic influence on the compost and the ambiguity of the origin of *A. baumannii* throughout this setting, we next attempted to identify soil habitats in more pristine environments. Searching for wet habitats rich in decomposing material where earthworms can thrive, we identified a site at the bank of the river Holtemme near Wernigerode, Germany, where we successfully isolated *A. baumannii* from earthworms, soil samples and the rhizosphere of different plants (*Urtica* spp., *Impatiens* spp., different grass species etc.). This site at the river bank with an area of only approx. 0.1 m^2^ turned out to be a “hot spot” for the isolation of *A. baumannii* and yielded 22 isolates within 3 months representing 18 different variants of OXA-51 (Suppl. Fig. S11). Moreover, samples collected at this specific site yielded 10 isolates on a single day, each harbouring a different OXA-51 variant. Strikingly, this site has remained a “hot spot” over the years even though the river bank has been remodeled several times due to flooding. None of the many additional sites we repeatedly sampled over the years along the river Holtemme showed any comparably high isolation rate.

Concomitantly, we had collected soil samples from a forest area south of Wernigerode, representing both deciduous and coniferous forests of various compositions at an altitude between 300 and 450 m above sea level. Again, we chose wet habitats with decomposing plant material, mostly alongside creeks and ponds, however, of 269 samples only two were positive for *A. baumannii* (0.7%) (Suppl. Fig. S12). To prove the deduced principle that *A. baumannii* preferentially colonize alongside creeks but not within forest areas, we selected a creek not previously sampled which has its source within the forest and which, after leaving the forest, flows through grassland before reaching the first village. We were unable to isolate *A. baumannii* from any sample collected inside or along the edge of the forest although the creek’s banks were rich in decomposing material. The first positive sample was instead collected at a distance of approx. 300 m to the edge of the woods, but positive samples remained rare until the creek had passed the first village (Suppl. Fig. S13). After passage of this village, we discovered two “hot spots”, which repeatedly collected positive samples (6/12 positive samples (50%) and 7/12 positive samples (58%), respectively). Collectively, forests do not offer soil habitats supportive of *A. baumannii*, not even alongside creeks, whereas *A. baumannii* can thrive alongside creeks and rivers outside of forests even if they are lined with trees. There are “hot spots” of *A. baumannii* where they can occur in striking diversity outside forests.

### Diversity in terms of OXA-51 variants collected from Poland and Germany

Currently, there exist 380 assigned variants of the β-lactamase OXA-51 protein encoded by the intrinsic *bla*_OXA-51-like_ gene of *A. baumannii* ((79) http://www.bldb.eu/, accessed October 31^st^, 2023). Our collection of strains encompasses 209 variants, thus about 55% of the presently known worldwide diversity. Of the 209 variants representing our collection, 125 were previously undescribed variants that were deposited in the course of this study (Suppl. Table S3). Given that some of these OXA-variants have been shown to be useful as indicators of phylogenetic relationships, our findings suggested that our collection might represent a considerable portion of the worldwide known diversity of lineages. To substantiate this hypothesis, we performed extensive sequencing and phylogenomic analyses.

### Phylogenomics-based representation of the diversity of our collection

From our collection of more than 1,450 non-redundant *A. baumannii* isolates collected outside the hospital context (Suppl. Table S1), we chose at least one strain representing each OXA-variant for whole genome sequencing. Further, we selected few OXA-variants for the sequencing of larger sets of representative strains, in order to illustrate the diversity within these supposed lineages (e.g. OXA-104, −106, −126, 343, −374, −378, −431), and in addition, selected several strains representing each of the OXA-variants associated with the international clones 1 to 8 (OXA-51, −64, −65, −66, −68, −69, −71, −90) (135). Moreover, due to the higher diversity of carbapenem-susceptible clinical isolates compared to carbapenem-resistant ones (102), 15 carbapenem-susceptible human clinical isolates from Germany were selected according to their encoding OXA-variant, some of the previously described (126). Finally, the collection was supplemented by a diverse set of 21 isolates covering veterinarian und human clinical isolates from a previous study (77). Altogether, our study provides 401 novel genomes, of which 229 originated from white stork nestling isolates, 77 from soil and plant root samples, 2 from air samples above compost, 59 from diverse animal materials including earthworms, rodents and shrews, pellets from diverse bird species, egg shells, feathers and nesting materials, 7 from veterinarian samples, and 27 from human clinical samples. For comparison to the previously available diversity within the species *A. baumannii*, a generic protocol selected 413 additional genomes available within public databases (Suppl. Table S4).

The pan-genome of this collection includes 50,989 unique genes, given a threshold of 90% sequence identity on the protein level (Table 1). The phylogenomic tree based on the core genome set of 1,728 conserved genes representing 1.328 megabasepairs (Mbp) of *A. baumannii* genomic sequences is illustrated in Fig. 3. Especially remarkable is the share of deeply branching lineages, suggesting massive radiation early in evolution of the species. Most of the isolates differ by 18,000 to 23,000 single nucleotide polymorphisms (SNPs) between each other (Suppl. Table S5). While only a few OXA-variants represented multiple times in this study are apparently monophyletic, such as OXA−431 or OXA−68, most OXA-variants show phylogenetic clustering (see interactive microreact project at https://microreact.org/project/3ApuGKD61qPLT1ZNmoKcTb-acinetobacternobaps12dec23 and Suppl. Fig. S14). In contrast, few OXA-variants show broad scattering, in particular the prototypic OXA-51 and some other OXA-variants (OXA−65, OXA−69, OXA−71) found associated with international clones (ICs) (135) (Suppl. Fig. S14). Another remarkable observation is that the collection of wildlife and environmental strains from Poland (n=235) and Germany (n=159) respectively, each covers a considerable portion of the worldwide known diversity (Fig. 4), pointing to an efficient mechanism of global spread. Moreover, human, animal and environmental isolates likewise spread all-over the phylogenetic tree (Fig. 5). Of particular interest are the many clades represented by environmental, animal and human clinical isolates, underpinning the relevancy of the One-Health concept in regards to *A. baumannii*. This also includes representatives of international clones, namely IC8 (Suppl. Fig. S15). Additionally, IC4, IC5, IC6 and IC7 are now also represented by at least one isolate from wild animals, livestock or the environment. Distances between human clinical isolates and their related isolates from environmental or wildlife sources cover a broad range, approaching distances of 27 SNPs within the core genome in the case of a human clinical isolate from Canada related to a white stork isolate from Poland (both OXA−64), 35 SNPs between a white stork isolate from Poland and a human clinical isolate from Thailand (both OXA−433), or 48 SNPs between a human clinical isolate from Japan and another white stork isolate from Poland (both OXA−762) (Suppl. Tab. S4-S6). These results called for a molecular clock analysis to allow for temporal placement of the genetic and geographical distances. Accordingly, we performed BEAST analyses (43) on the OXA−126 clade (n=34), revealing a mutation rate of 1.1345 x 10^−6^ (+/− 3.2 x 10^−7^) per site per year corresponding to approx. 1.5 SNPs per year within the core genome (range 1,08−1,93 SNPs per year), slightly lower than the 1.5 substitutions per site per year reported for IC1 (43) (Suppl. Tab. S7). The most distant isolates within the OXA−126 clade, 13-291−1C and 19-Pos71−1, differ by 12,354 SNPs corresponding to 8,236 years (range ∼6,400−11,440 years), pointing to a strong and constant selection pressure stabilizing the OXA−126 variant within the natural context. Applying this molecular clock on the species level, the most recent common ancestor of all members of the species *A. baumannii* likely existed only around 15,000 years ago. Accordingly when applied to the closest relatives found between clinical and environmental/wildlife isolates in our study, those intercontinental pairs of isolates mentioned above share their most recent common ancestor only 18−32 years before present. Further relationships of particular interest are listed in Suppl. Table S6.

**Fig. 3:**
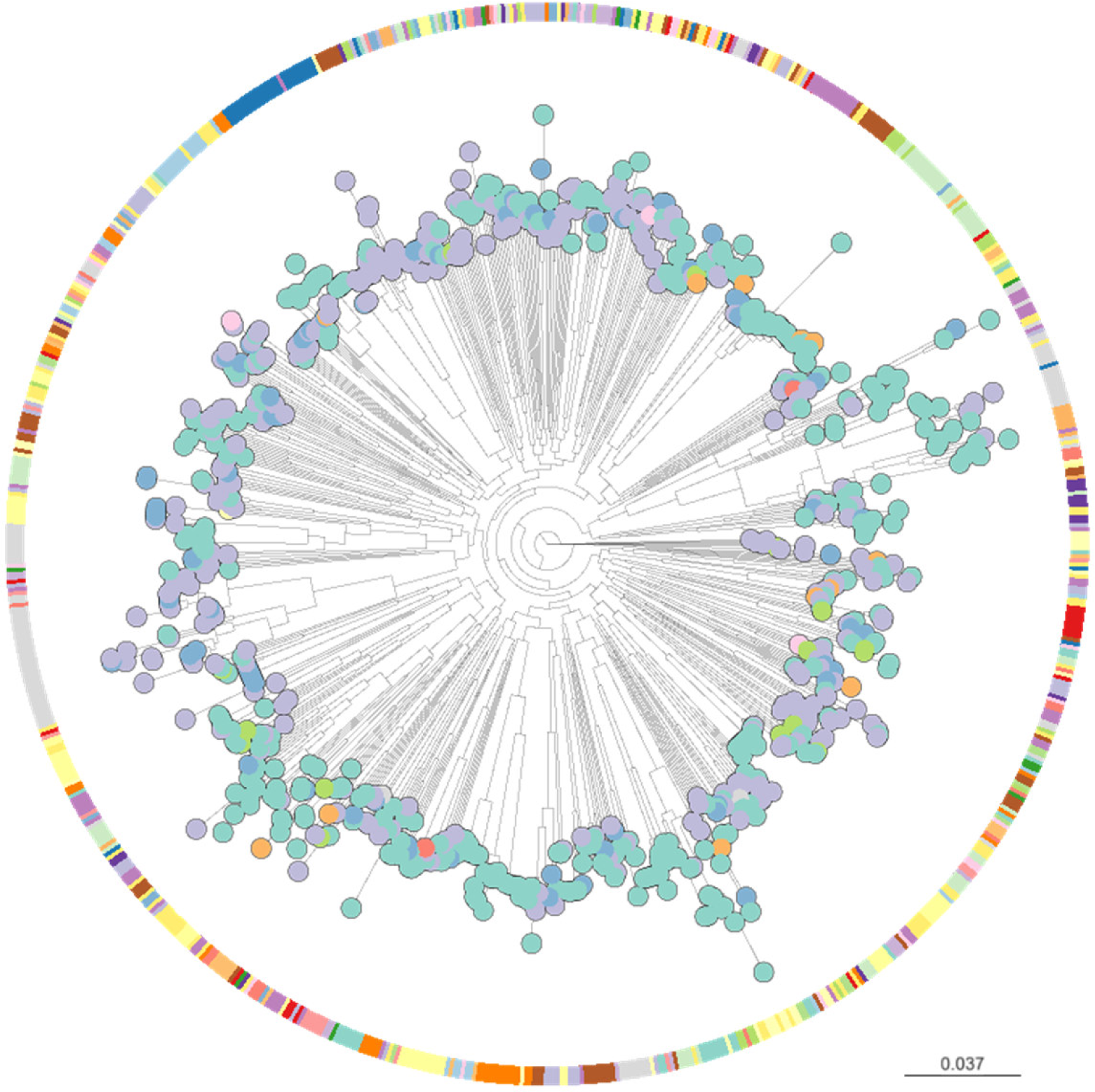
Phylogenomic tree of *A. baumannii* based on the core genome of a set of 826 genomes. Colors by OXA-51 variant types. See microreact project at https://microreact.org/project/3ApuGKD61qPLT1ZNmoKcTb-acinetobacternobaps12dec23 for further analysis and illustration of the dataset.

**Fig. 4:**
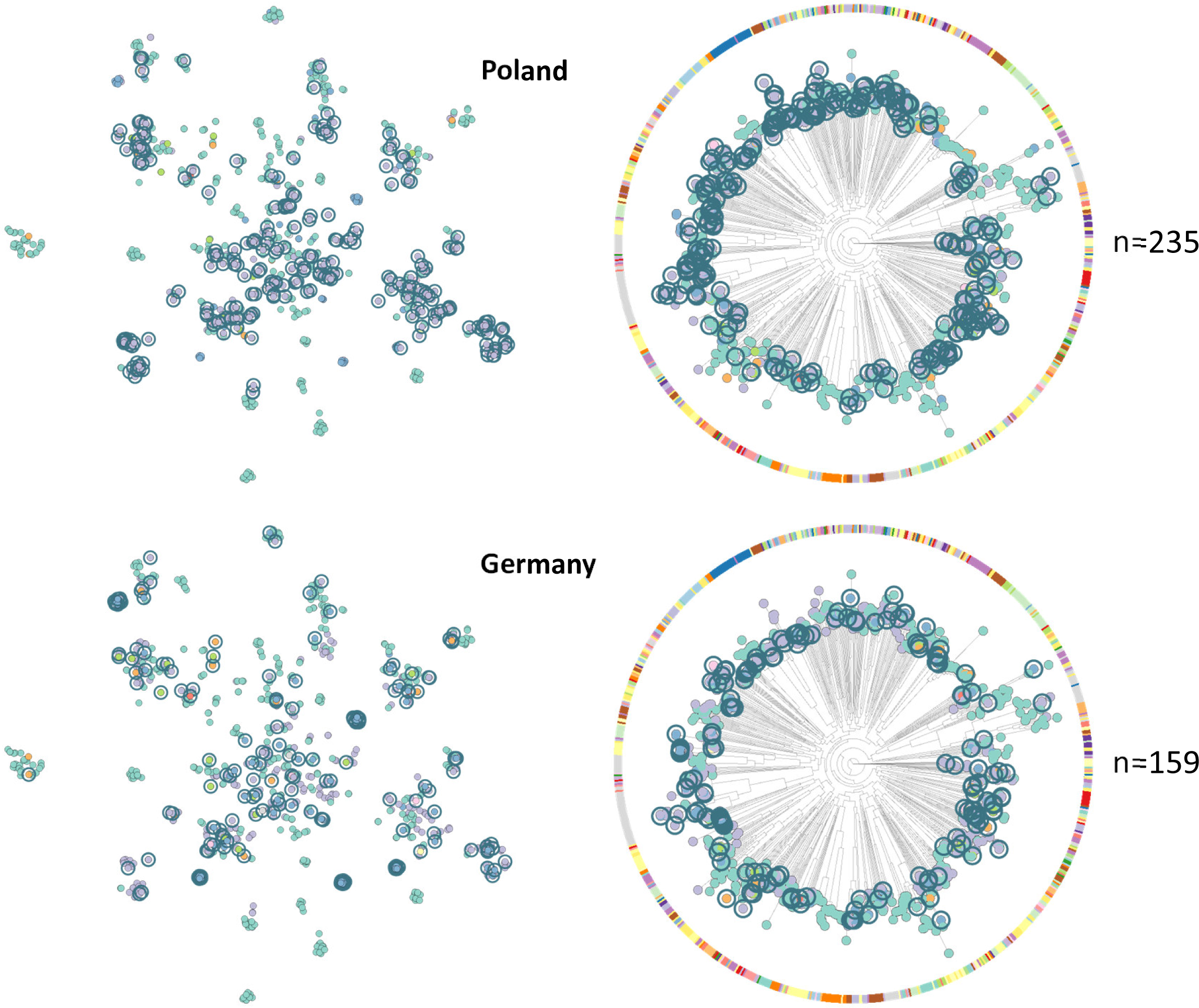
Isolates from Poland and Germany represent a significant part of the global diversity. Isolates from Poland and Germany, respectively, indicated by grey blue circles. See microreact project at https://microreact.org/project/3ApuGKD61qPLT1ZNmoKcTb-acinetobacternobaps12dec23 for further analysis and illustration of the dataset; perplexity clustering to the left (p=10).

**Fig. 5:**
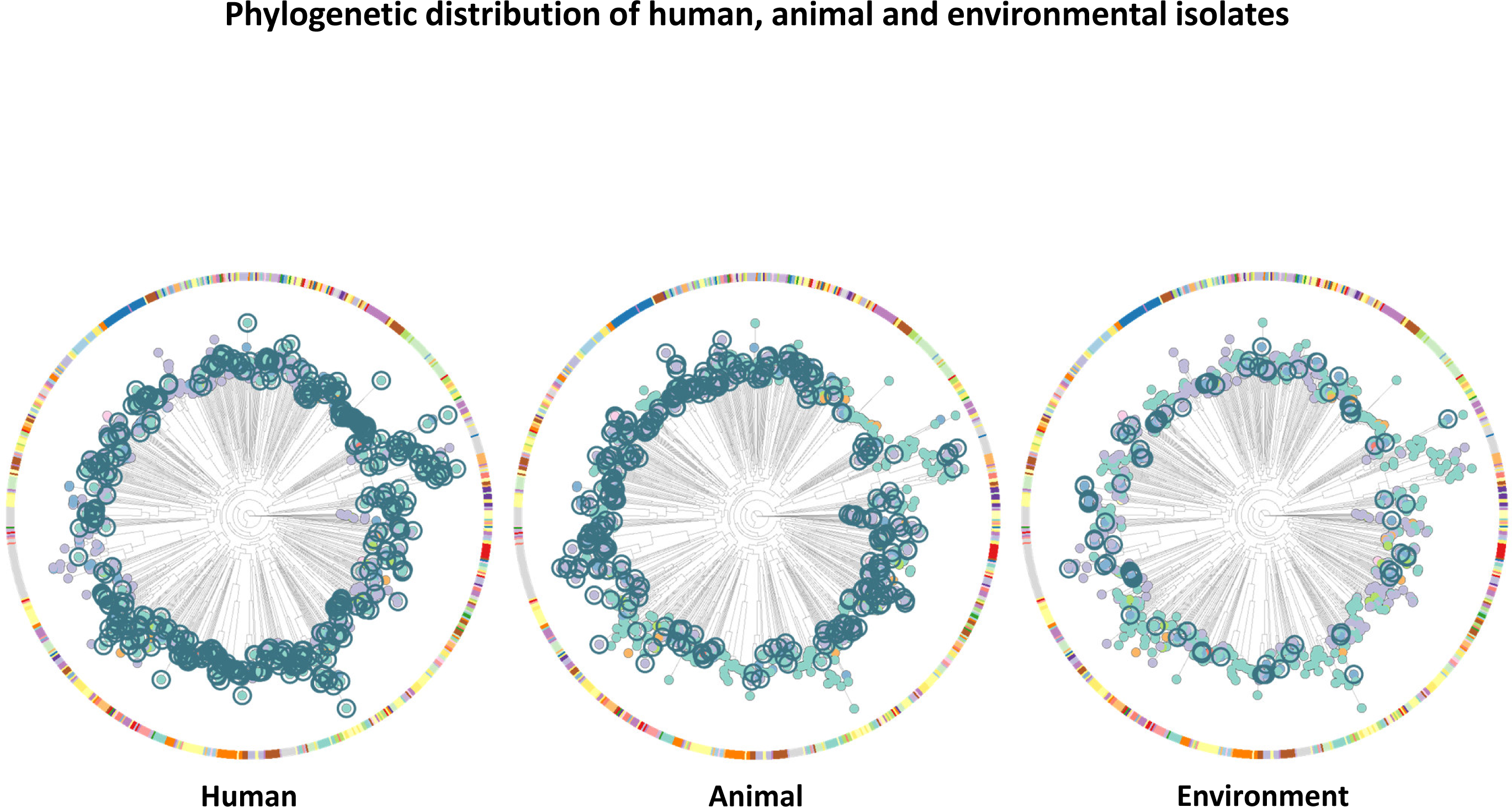
One-Health perspective: Human, animal and environmental samples are distributed all over the tree. Isolates of human, animal and environmental origin indicated by grey blue circles. See microreact project at https://microreact.org/project/3ApuGKD61qPLT1ZNmoKcTb-acinetobacternobaps12dec23 for further analysis and illustration of the dataset.

**Table 1:**
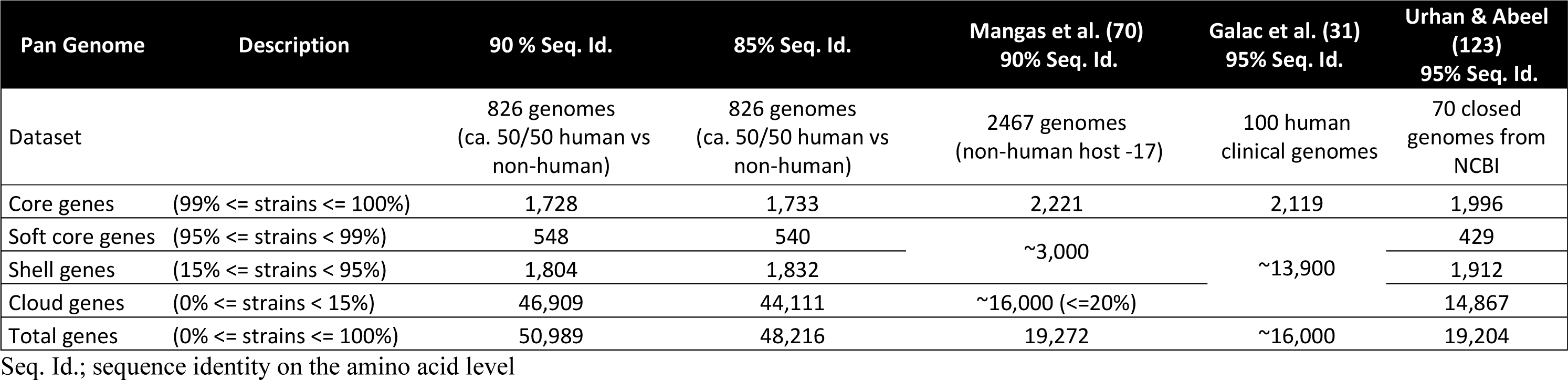
Pan-genome analysis: comparison with previous studies.

### Natural endowment with resistance genes and IS elements

Altogether, intercontinental spread between North America, Europe and Asia within a few years cannot be explained by the migratory activity of white storks which migrate between Europe and Africa. Rather, human activity, or natural mechanisms of airborne spread, need to be considered. Hence, we analyzed the genomes regarding their endowment with antibiotic resistance genes and insertion sequences (IS elements) as indicators of anthropogenic selection pressure (Suppl. Tab. S4; Suppl. Fig. S16-S17). Globally, the newly sequenced isolates harbour an intrinsic *bla*_OXA-51-like_ gene (96% of the collection), a majority of which also harbour the *bla*ADC gene in one of 43 variants. Nickel resistance gene *nreB* and efflux pump gene *amvA* are also highly abundant. Moreover, most isolates also harbour the *ant(3’’)-IIa* aminoglycoside and the *abaF* fosfomycin resistance genes, suggesting these also to be part of the intrinsic genomic endowment of the species. Additionally acquired resistance genes are limited to a few isolates, mostly collected with a veterinary or human clinical contexts, or within the context of livestock production. Environmental and wildlife-associated isolates do not harbour these acquired resistance genes known from the clinical isolates indicating the comparably pristine habitats they originate from. Similarly, while IS elements are rarely enriched in environmental and wildlife-associated isolates, clinically related strains harbour an increased amount of IS elements (Suppl. Table S4, Suppl. Fig. S17). Moreover, few of our environmental or wild animal isolates harbored an interrupted *comM* gene, a configuration typically found in multidrug-resistant lineages firmly established in the hospital (36). Altogether, this suggests that our environmental and wildlife-associated isolates show little signs of anthropogenic selection pressure and indeed originate from pristine environments.

### Colonization of compost material and linkages to the fungal world

As our phylogenomic analyses suggested global spread of clonal lineages long before globalization of modern times, we considered various modes of dispersion independent of human activity. Our samplings in garden compost suggest that colonization of *A. baumannii* is associated with decomposition of fresh plant material, as *A. baumannii* vanishes after the feed of the compost is stopped. This points to attraction of *A. baumannii* throughout the early steps of decomposition, during which fungi are key players. We thus reasoned that *A. baumannii* might start colonization together with fungal spores that could serve as a vehicle for local as well as global spread of *A. baumannii*. To prove an airborne spread of *A. baumannii* in a local setting we mounted open tubes in a distance of 50 cm above an active compost with the opening positioned downwards or the tube positioned vertically for 24 hours. Then the tubes were closed, transported to the laboratory, filled with liquid medium and processed to cultivate *A. baumannii*. As this *ad hoc* setting instantly yielded several isolates of *A. baumannii*, we next designed experiments to study the colonization of sterilized fresh plant material deposited in the garden. The autoclaved plant material was deposited either directly in the garden in a distance of 5 to 10 m from the compost, or it was deposited in a sterilized plastic bag either open top and shielded from the soil or shielded from the top and accessible from the soil. We found that every setting was colonized within 2−3 weeks, indicating that colonization can occur not only via soil contact but also via the air.

Next, we tested if *A. baumannii* is capable of attaching to fungal spores using *Aspergillus* spp., which are known to play a key role during the first steps of plant decomposition and dominate within the compost setting (38). We incubated spores from different *Aspergillus* spp. isolates with different strains of *A. baumannii* and observed a marked adherence of the bacteria to fungal spores (Fig. 6). Similarly, adherence to spores from different *Penicillium* spp. isolates was observed. Interestingly, adherence was not an *ad hoc* phenomenon but required 2 to 3 hours of incubation to manifest and resulted in an inhibition of spore germination, a process that typically became observable after 5−6 hours of incubation in our setting. Taken together, these experiments demonstrate that *A. baumannii* has the capacity to interact with fungal spores and to colonize new habitats via the air.

**Fig. 6:**
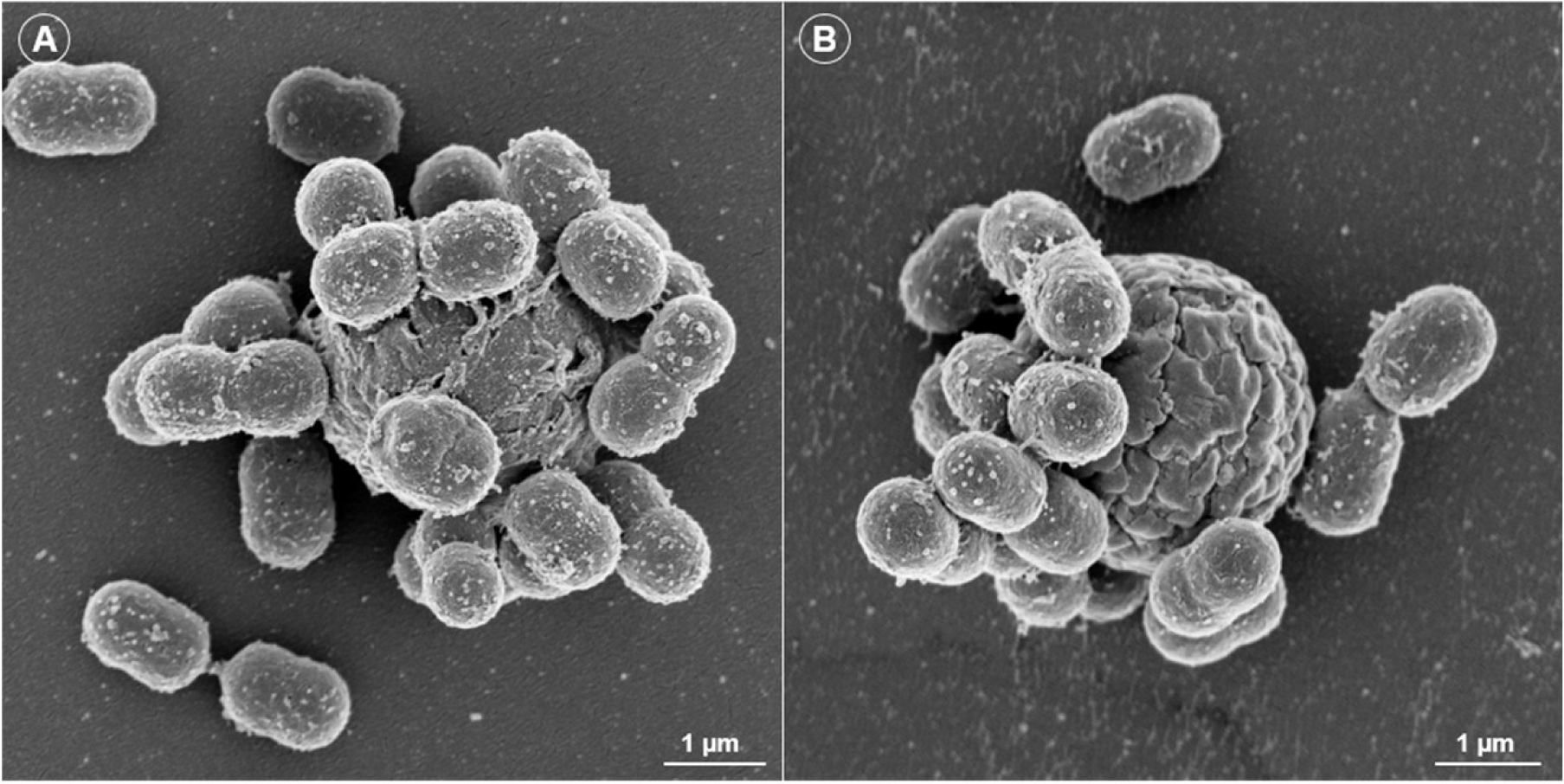
Scanning electron microscopy on *A. baumannii* adhering to *Aspergillus* spores. (A) *A. baumannii* 31D1 adhering to a spore of *Aspergillus niger* complex isolate U17-Zw-P20. (B) *A. baumannii* 31D1 adhering to a spore of *Aspergillus quadrilineatus* strain Eld3.

### The “where” and “when” of sampling

Finally, we challenged our current model of the prevalence of *A. baumannii* in natural habitats. We chose to collect samples during late summer when positive rates at our hotspot sites peaked. We collected 20 samples from river banks within forests and 20 samples from river banks outside of forests across the Harz district, Germany. For both groups we chose sites we had never sampled previously and with indicators of nutrient-rich soil, such as deposition of decomposing plant material and growth of nettles (*Urtica* spp.), *Impatiens* spp. or *Aegopodium podagraria*. While 1 out of 20 (5%) forest samples was positive for *A. baumannii*, 6 out of 20 (30%) samples were positive from river banks outside of forests. The positive rate of 30% in selected soil samples thus is in the same range as that yielded from choana sampling of white stork nestlings.

## Discussion

Here, we present the most comprehensive study on the ecology of *A. baumannii* to date. Today, representatives of this species can be isolated from specific natural habitats at an unprecedented scale, and we hope that this will stimulate the scientific community to collect and characterize *A. baumannii* from natural habitats worldwide. Although our sample setting, essentially restricted to samples from Poland and Germany, represents a significant portion of the worldwide known diversity, evidently, our picture remains incomplete. In particular, we have not yet collected representatives of the international clones IC1 - IC3 from natural environments, including the clinically dominant IC2, suggesting that their relatives either prefer completely different habitats or are geographically restricted. We focus the discussion on the interpretation of our data in light of understanding the evolution behind this species and in particular adaptation to the hospital environment.

### Comparison of the *A. baumannii* population structure to *Pseudomonas aeruginosa*

It is interesting to compare the population structure of *A. baumannii* with that of another cosmopolitan opportunistic pathogen within the order of *Pseudomonadales*, *Pseudomonas aeruginosa*. Likewise, *A. baumannii* and *P. aeruginosa* are found worldwide across various habitats, including soil. Similarly, a local study setting also captured a significant fraction of the global diversity within the species *P. aeruginosa* (93, 100). Possibly, the two species share similar modes of global spread and future studies should compare their velocities of spread to challenge this hypothesis (67).

However, in striking contrast to *A. baumannii*, the global diversity of *P. aeruginosa* consists of only five groups, two of which dominate the scene both as human opportunistic pathogens and environmental bacteria (27). In *P. aeruginosa*, the dominant clones characterized within the context of human infection are also found in the environment, albeit at a skewed abundance (127), while the dominant clone of *A. baumannii* in hospitals (IC2) has not been isolated from pristine environments in our study or elsewhere. It is also interesting to note that the pan-genome of *A. baumannii* determined in our study is in the same order of magnitude as that of *P. aeruginosa* (50,989 versus 54,272), while the core gene sets consist of 1,728 and 665 genes, respectively, and the average genome size of *A. baumannii* is also considerably smaller compared to that of *P. aeruginosa* (4 Mbp versus 6−7 Mbp) (27).

Altogether, these findings not only indicate a significantly different evolution of both species in the environment but also differently selecting bottlenecks for entry and establishment of environmental isolates in the hospital setting.

### Evidence for interrelatedness of *A. baumannii* life cycle with the fungal world

The linkages between the worlds of fungi and *A. baumannii* found in the literature are manifold and require a review for full appraisal. To justify our claims, we will discuss selected aspects and provide a more detailed list of published evidence in the Supplements (Suppl. Table S8). Tan et al. (116) found airway colonization with *Candida* spp. yeasts to be an independent risk factor for *A. baumannii* ventilator-associated pneumonia. This finding was consolidated in an infection model of the rat lung (115). In a pioneering paper, Smith et al. (108) described the synergistic interaction between the yeast *Saccharomyces cerevisiae* and *A. baumannii*, with ethanol produced by the fungi stimulating the virulence of *A. baumannii* in the nematode host *Caenorhabditis elegans*. Seminal work by Peleg et al. (90, 91) applying a co-infection model of *A. baumannii* together with the yeast *Candida albicans* in *C. elegans* revealed a complex interplay between bacteria and fungi. While *A. baumannii* suppressed filamentation of the fungus, the latter arranged counteroffensive via the quorum sensing molecule farnesol. Farnesol was later demonstrated to disrupt cell membrane integrity in *A. baumannii* and to impair biofilm formation and motility (59). The virulence potential of *A. baumannii* against *C. albicans* has been further characterized by others (30, 74, 78). Moreover, several isolates of *A. baumannii* have been described to suppress phytopathogenic fungi (64, 113). The microbiome analysis of household dust revealed a significant positive correlation of *Acinetobacter* with several fungal genera such as *Alternaria*, *Aspergillus* and *Fusarium* (20). Further, it is worthwhile to note that 1,3-diaminopropane, the dominant polyamine of *Acinetobacter* with a known role in virulence of *A. baumannii* (107), upregulates secondary metabolism in fungi such as *Aspergillus* and *Penicillium*, including the biosynthesis of β-lactams (73, 139). In this light, we speculate that as a result of co-evolution with fungi producing differing β-lactam variants, a multitude of OXA-51 variants have evolved in *A. baumannii*.

*Aspergillus* is a major cause of morbidity and mortality in a wide range of birds including specific lineages of poultry (117) and particularly so in white stork nestlings (86). It is important to keep in mind that azole antifungal drugs are not only used in the hospital setting but also in agriculture (15). This may influence the *Aspergillus*-*Acinetobacter* relationship with a possible impact on the abundance of *A. baumannii*. Consistently, our data indicate that *A. baumannii* prevalence in white stork samples is positively correlated with heterogeneous agricultural use (Suppl. Fig. S18 and Suppl. Material S1).

Sporadically, *A. baumannii* infection has been reported following *Aspergillus* infection in humans (69), but metagenomics-based analyses suggest that fungal and bacterial co-infections of the lung are common, with *A. baumannii* being one of the dominant pathogens (138). Concerning our hypothesis that *A. baumannii* could use *Aspergillus* and other fungal spores for hitchhiking, it is important to emphasize that both *Acinetobacter* and *Aspergillus* readily aerosolize (38). In the light of all this groundwork, our findings about the interaction between fungal spores and *A. baumannii* are far from surprising. It is however important to consider these linkages in the context of hygiene and nosocomial infections. For instance, mold growth on moist surfaces, a common problem in tropical regions, may also contribute to the spread of *Acinetobacter* in the hospital. Municipal composting plants should be also assessed carefully (38).

### Forest as a ‘no-go’ area for *A. baumannii*

Our data indicate that forests do not provide supportive habitats for *A. baumannii*. Particularly, we documented low isolation rates from forest soil samples and from samples collected in the vicinity of forests, and we found no association of *A. baumannii* with sylvan black storks. Measured pH values from soils supportive of *A. baumannii* were not significantly different from those of forest soil samples (data not shown). We therefore propose that *A. baumannii* is kept away from forests either via direct inhibition of *A. baumannii* by the fungi of the forests’ mycorrhizae or via suppression of those fungi supportive of *A. baumannii* colonization. In line with the latter hypothesis, fungi of the division *Basidiomycota* for example are known to produce antifungal compounds (61). If the former explanation should hold, forest microbiota should be intensively screened for potential antibiotic producers. Interestingly, a study from China on microbial emission levels depending on land use did not reveal a negative effect of forests on *Acinetobacter* abundance (62). Possibly, the situation in Europe differs from that in China or the general trend for the genus *Acinetobacter* does not represent the situation for *A. baumannii*. In line with the latter explanation, several *Acinetobacter* species such as *A. silvestris* (83) and *A. bohemicus* (60) have been isolated from forests. Although the microbial communities of forest soils are largely defined by pH (52) and dominant tree species (6), we could not observe a correlation of either of the two parameters with the lack of *A. baumannii* in forests (data not shown). It is evident that metagenomics data already available from forest soils need to be evaluated and complemented by new studies specifically involving *A. baumannii* habitats, such as those uncovered within this study.

### Early adaptive radiation within the species *A. baumannii* associated with Neolithic?

Our data suggest a massive radiation within the species *A. baumannii* early after its emergence. The average distance of the numerous unrelated lineages within the species is approx. 20,000 SNPs per genome based on our core set of genes (Suppl. Table S5). According to our molecular clock analyses and those of others (Holt et al., 2016), the substitution rates are in the range of 1.13 - 1.5 x 10^−6^ substitutions per site and per year. Based on our core genome alignment, this corresponds to 1.5 SNPs per core genome per year. Accordingly, most of the lineages are about 13,000 years apart from each other (range ∼10,400−18,500 years). This is in agreement with global warming during the beginning of the Holocene and early Neolithic when human activities started to change the environment producing a plethora of novel types of habitats, especially due to deforestation and later the development of agriculture and livestock farming. In this context, deforestation is of specific importance given the adverse impact of forests on *A. baumannii* as outlined above. In the European Alps, deforestation on small areas due to human activity has been dated back to about 10,000 years before present (32). Moreover, livestock farming may have additionally contributed to this radiation, given the association of *A. baumannii* with cattle, sheep, and poultry (56, 63, 129). All in all, our data suggest that human activity during the beginning of the Holocene and early Neolithic was a key driver of the adaptive radiation within the species. This possibly also applies to radiation within the *A. calcoaceticus*-*A. baumannii* complex in the earliest phase of the Neolithic, in line with the close relationship of species within this clade (21).

### *A. baumannii* – jet stream rider

Our local sample settings in Poland and Germany revealed a significant proportion of the worldwide known diversity of *A. baumannii*, with evolutionary distances to hitherto known genomes, mostly of clinical origin, ranging from very ancient to very recent. Thus, our data illustrate a massive global dispersal on the sub-species level since ancient times, which is consistent with the findings of Louca (67) that most prokaryotes are globally spread even at sub-species resolution. However, the velocity of spread even for human-associated prokaryotes exhibiting the highest diffusivity, was determined at only 580 kilometers per 100 years for individual lineages while for terrestrial bacteria it was only 370 km per 100 years. Intercontinental transmission rates of human-associated bacteria on a per-lineage basis were estimated at around 5.5 events per 10^6^ years for the most probable transfer between North America and Europe, i.e. one intercontinental transfer event in every ∼182,000 years (67). Evidently, the massive intercontinental spread suggested by our data is exceeding estimates for those most transmissible species, namely human-associated ones (67), by far. If modern human activity would account for the spread observed, a strong association of *A. baumannii* with the human microbiota could possibly explain the data, however such an association has been refuted (9). Alternatively, a strong association of *A. baumannii* with global flow of commodity, in particular livestock and meat which can be heavily contaminated with *A. baumannii* (68), might explain the global spread. However, as outlined above, the origin of our environmental and wildlife isolates from pristine environments is evident from their genomic structure. All in all, there is no evidence that the observed global pattern of distribution of environmental *A. baumannii* is caused by human activity. It is further not explainable by migratory patterns of white storks, which do not cross oceans (26). Based on our data and published work, we propose airborne spread as an original strategy of the species and possibly of other members of the genus *Acinetobacter*, likely facilitated by hitchhiking on fungal spores. In the hospital setting and livestock production facilities, airborne spread of *A. baumannii* has been already demonstrated (72, 132). The prerequisites for an effective long-distance atmospheric spread are tolerance to desiccation and radiation, as well as potential for aerosolization. The tolerance of *A. baumannii* to desiccation is well documented (47, 57, 88, 136). Further, resistance to radiation is not only described for *A. radioresistens*, but also for other *Acinetobacter* species, including *A. baumannii* (16). Moreover, *Acinetobacter* spp. are among the dominant airborne bacteria found in bioaerosols (62, 76). The aerosolization behavior of bacteria and fungi from vegetable waste compost has been studied in detail, revealing efficient aerosolization of both *Acinetobacter* and various fungi including *Aspergillus* (38). In line with these findings, *Acinetobacter* is abundant in rainwater (1, 5). What is more, glacier microbiomes show the highest relative abundance of *A. baumannii* and *A. junii* compared to all other potential pathogens (65). Taken together, there is evidence illustrating that *A. baumannii* is well adapted to survive atmospheric long-distance spread. This scenario may apply to other species of the genus *Acinetobacter* but also to other bacteria with linkages to the fungal world such as other members of the *Pseudomonadales* (90).

After future increase of studies focusing on environmental, animal and non-MDR isolates and their corresponding genomes from all-over the globe, it should be possible to challenge (67) the “jet stream rider” hypothesis. It would claim that because of jet streams‘ west to east streaming and their hemispheric association, spread between e.g. North America and Europe should be more likely than between North and South America. In line with this hypothesis, the diversity of clinical isolates from South America differs significantly from those observed in the northern hemisphere (85), but at present ecological differences cannot be excluded as a reason for this phenomenon.

### Seasonality and entry into the hospital environment

Seasonality of *Acinetobacter* infections with a global peak in summer has been discussed (97). However, seasonality has not been observed for MDR *Acinetobacter* (28), which show a high degree of clonality and are known to be transmitted from patient to patient and from hospital to hospital in contrast to the diverse group of non-MDR *Acinetobacter* (102, 126). Our data on the occurrence of *A. baumannii* in compost and soil samples as well as birds’ pellets and small mammals approve seasonality with a summer peak. In line, association of *A. baumannii* with cattle also exhibits seasonality, peaking between May and August (56). Noteworthy, aerosolization of bacteria and fungi also peaks in summer (131), as discussed above. In conclusion, we provide further evidence that preferentially during the summer season, when *A. baumannii* thrive in natural habitats, novel lineages of *A. baumannii* enter the hospital environment worldwide causing increased infection rates with non-MDR appearance.

### Pan-genome and potential for further development of the species

Previous analyses of the pan-genome of the species *A. baumannii* ranged from 16,000-20,000 genes (31, 70, 123). Here, we present a conservative estimate of 50,989 different genes, more than doubling previous appraisals (Table 1). This drastic increase in the size of the pan-genome of the species illustrates the significant bias due to the focus on MDR isolates (the majority of which belong to a few widely disseminated clonal lineages) within the scientific community, but also indicates that compared to the broad diversity of lineages in nature, only few have managed to establish themselves within the hospital environment. Our data set will now facilitate identification of the critical factors of success owned by international clones. Even though a few established lineages dominate, there is a constant invasion of novel lineages into the hospital, particularly in summer. Given the enormous genomic diversity and fluidity illustrated here, combined with the capacity to recombine excessively (34, 130), a huge potential is evident for further development including the adaptation to the hospital environment and humans as hosts. This potential must not be underestimated given the recent development of a hypervirulent sub-lineage of IC2 isolated from sheep (63). Note that as *A. baumannii* is apparently a very recent species, with its radiation starting only around 15,000 years ago, its adaptation to humans and livestock might be only in its beginnings. Moreover, we also need to consider the potential interspecies transfer of DNA from other nosocomial pathogens (120).

### *A. baumannii* – opportunist by nature

We have demonstrated that sterilized organic matter deposited under the open sky is colonized by *A. baumannii* within 2−3 weeks during summer. Our data suggest that *A. baumannii* is present in various environmental habitats and in the air, albeit at generally low abundance and with marked seasonality. Consequently, almost all humans have already been exposed to *A. baumannii*. However, only few people suffering from predisposing diseases, mostly in tropical regions, face community-acquired infections from *A. baumannii* (18) suggesting a very low virulence potential and/or high infection doses required to establish an infection. It is only in the hospital context that *A. baumannii* becomes a more pronounced isssue, even more in intensive care units (ICUs) and in tropical countries. It appears to be the nature of this species to patrol the environment in search of conducive habitats and it is our challenge to protect the most vulnerable from this opportunistic contact.

The transient nature of *A. baumannii* occurrence is not only due to seasonality but also due to succession in its habitats. So far there is no evidence of permanent colonization of any habitat with *A. baumannii*. If compost feeding is stopped, *A. baumannii* vanishes. Likewise, the probability of isolating *A. baumannii* from white stork nestlings decreases with the age of the chicks (Suppl. Fig. S19, Suppl. Material S1). Similarly, samples from commercially reared turkey chicks are heavily loaded with *A. baumannii* at the first day of life, but this load vanishes during life (103). Where permanent establishment and local spread is not possible, patrolling in the air appears as an efficient strategy to identify the next opportunity for colonization. Several studies resolved on the level of the genus *Acinetobacter* illustrate the marked position of *Acinetobacter* species at the beginning of a succession. Aging of cattle and chicken manure correlates with a characteristic burst and decline of *Acinetobacter* (17, 84, 92). In line, *Acinetobacter* has been described to dominate the rice paddy rhizosphere after green manure treatment in comparison to winter fallow treatment (137). Taken together, to be in the right place at the right time is the challenge valid for both the bug and the bug hunter.

### One-Health perspective

Given the broad distribution of human clinical isolates, animal and environmental isolates all-over the phylogenetic tree and the linkages between isolates of different sources disclosed, the capability to cause opportunistic infections appears as an ancient skill of the species rather than that of a few recent human-adapted lineages. In line with this, previous studies on avian isolates did not reveal significant differences in virulence compared to human clinical isolates (129). Studies on wastewater discharge and river water indicate spreading of clinical isolates of *A. baumannii* into the environment in Poland and elsewhere (44, 51, 105). However, the IS*Aba*1/*bla*_OXA-51-like_ genetic configuration observed in carbapenem-resistant *A. baumannii* from wastewater and river water in Poland by Hubeny et al. (44) has not been observed in any of our environmental and wildlife isolates indicating their still pristine context. Interestingly, the same is true for the compost setting studied here indicating that the source of compost isolates is from nature rather than from the anthropogenic context. Thus, our sample and data collection can serve to represent the baseline of nativeness to study the evolution of lineages towards adaptation to the hospital environment and multidrug resistance. It will also help to decipher the role of livestock farming as a potential accelerant to the development of some lineages. We emphatically support the call to action recently expressed by others to implement a One-Health perspective on *A. baumannii* (14, 40, 124). This should not only include pet, livestock and wildlife animals as well as environmental sampling, but also genome-based studying of non-MDR clinical isolates as these presumably only recently entered the hospital setting.

### Miscellaneous

Our findings outline that *A. baumannii* can be effectively isolated from wet and nutrient-rich soil alongside waters, which is in line with previous studies (3, 35, 55, 96, 112). The natural resistance endowment of *A. baumannii* indicates evolution in an environment rich in antibiotic producers, such as the soil habitats verified here. Note, however, that also the adaptation to airborne spread is linked to antibiotic resistance. Resistance to UV radiation involves enzymes dedicated to the detoxification of reactive oxygen species (ROS) such as catalases and superoxide dismutases (101, 111), and these enzymes, ROS detoxification and thus redox homeostasis crucially determine deployment of antibiotic activity (2, 23, 39). Moreover, *Aspergillus* spores are coated with antimicrobial peptides (22). Adaptation to this specific niche therefore requires resistance mechanisms, which again may contribute to resistance development of *A. baumannii* in the hospital setting.

Stork nests are typically used for several years or even decades with a continuous input of organic matter resulting in soil formation and the settlement of soil organisms as well as plants (10, 24). Thus, a stork’s nest is the perfect reproduction of a compost-like habitat, so that fungi-*A. baumannii* aerosols from the nest should be considered as a potential source of *A. baumannii* colonizing nestlings. In line with this, nest material and egg shells from white storks revealed *A. baumannii* contamination rates of 30−40% (Suppl. Table S1). Future studies should correlate the age of nests with the probability of colonization of nestlings within these environments.

A recent study found no *Acinetobacter* in faeces collected from white storks in Spain (46). This is in accordance with the low abundance documented in material from Spanish storks here and more so since the samples for the cited study were collected during winter. Comparing our data on white stork nestlings from Poland and Spain, it is evident that there are significant ecological differences. Of 60 eggs tested in Spain not a single one was positive for *A. baumannii*, while 27 out of 68 eggs (40%) tested in Poland (voivodship Greater Poland) were contaminated with *A. baumannii*.

Our groundwork now offers the potential to study horizontal gene transfer (HGT) on-site. The diversity found at a specific site reached 20 distinct lineages during a season and up to 10 distinct lineages could be isolated from a single sample site on a single day (Suppl. Fig. S11). Extensive HGT was detected between co-colonizing lineages (Suppl. Fig. S20).

### Open questions

Among the most interesting questions is where and how *A. baumannii* survives during the winter season. Given its marked tolerance to desiccation stress, dormancy in dry places might be a simple explanation (57). Alternatively, there might be (micro-)habitats where they are permanently established and from where they spread again at opportunity. Soil-dwelling amoebae should be considered potential hosts (12, 114). Since colonization rates of earthworms were found lower than those of ambient soil we do not consider earthworms as original reservoir although they might contribute to spread. In this regard, it is interesting to note that other bird species partially feeding on earthworms such as the blackbird *Turdus merula* and the European robin *Erithacus rubecula* do not exhibit colonization with *A. baumannii* (66) once again pointing to a very specific ecological setting of the white stork nestling.

### Limitations

Reliable isolation of *A. baumannii* from clinical samples is challenging (71) and so is from environmental material (134). Recently, evidence was presented that CHROMagar Acinetobacter might introduce a bias in isolation so that specific lineages might not be represented (134). Accordingly, alternative enrichment and isolation protocols should be run in parallel to complete our picture. Although we could demonstrate worldwide spread of lineages isolated in Poland and Germany, not all lineages necessarily spread worldwide. Ecology of *A. baumannii* in other climates and geographic regions may differ considerably.

### Our current view on the ecology of *A. baumannii*

In a nutshell, our collective data predict that the primary habitats of *A. baumannii* are associated with soil. Decomposition of plant material by fungi appears to set the stage for proliferation of *A. baumannii*. After the first steps of decomposition they rapidly leave the scene and spread via aerosols, possibly including hitchhiking on fungal spores, to patrol in search of novel conducive habitats. This opportunistic lifestyle requires effective basic equipment to withstand antibiotics produced by fungi and other soil-dwelling organisms, as well as rapid adaptability to novel and changing environments which is facilitated via its notable potential to undergo horizontal gene transfer. More so, the capability to spread via the atmosphere on a global scale co-evolved with adaptation to desiccation and radiation stress. These traits are key to a successful establishment in the hospital environment. The virulence potential of this opportunistic pathogen is low, ancient and currently not specific to humans. However, in terms of evolutionary history the species is still young and has a tremendous potential to further develop into a human and animal pathogen. Key virulence factors such as iron siderophore and capsule biogenesis might have evolved to compete in the soil and to resist phagocytosis by amoebae, respectively. While the mechanism of global dispersal of *A. baumannii* in nature appears to be ancient and not a direct result of human activity, evolution of the species and present-day occurrence is significantly impacted by humans’ land use and its influences on habitats favorable to *A. baumannii*.

## Experimental Procedures

### Sample collection and processing

Agar gel medium transport swabs (COPAN 108C, HAIN Lifescience, Germany) were used for sampling of white stork nestlings as previously described (129), and COPAN 110C swabs were analogously used to sample dead rodents and shrew (tracheal and rectal sampling). Swabs were immediately transferred to Amies transport medium and stored at 4°C until direct plating on CHROMagar^TM^ Acinetobacter (CHROMagar, France). CHROMagar Acinetobacter was prepared according to the manufacturer’s description without addition of the CHROMagar MDR supplement CR102. Pellet and soil samples were preincubated in minimal salt medium supplemented with 0.2% acetate for 5 hours at 37°C as previously described (134) prior to plating on CHROMagar Acinetobacter.

### Garden compost

The compost was fed with plant remains from the garden, vegetable kitchen waste and egg shells which were deposited within a wooden frame (length/width/height: 0.8 m/0.8 m/0.5 m), the compost regime was as follows: feeding for twelve months beginning in spring (approx. March), afterwards rest period for 12 months without relocation and without stirring or turning, then spreading into garden, continuous sampling over the complete two years (approx. 10 cm below surface (sampling with metal shovel and disinfection wipes); occasionally, sampling was applied with a boring rod of an effective length of 1 m (Bodenprobentechnik Peters, Germany); samples were taken every 10 cm.

### Bacterial species identification

Species determination of isolates recovered from CHROMagar Acinetobacter was based on PCR detection of *bla*_OXA-51-like_ (122), partial 16S rRNA gene sequencing (125), and partial *rpoB* sequencing using primers Ac696F and Ac1598R as described previously (82). To determine *bla*_OXA-51-like_ variation the coding region was fully sequenced as described previously (135). New OXA-51 variants were assigned and deposited at GenΒank (Suppl. Table S3).

### Illumina sequencing

Libraries for Illumina short read sequencing were prepared from 1 ng of extracted DNA utilizing the Nextera XT DNA Library Prep Kit according to the manufacturer’s recommendations (Illumina Inc., USA). Sequencing was carried out in paired-end (2×300 base pairs) on a MiSeq benchtop instrument. Quality control included the removal of adapter sequences, minor contaminations and short contigs below 700 bp in length through an in-house pipeline. The whole genome shotgun project of 401 isolates has been deposited at GenBank under the BioProject accession PRJNA862736.

### Genomic reconstruction and gene-marker analysis

Short-read DNA fragments were successively utilized to re-construct high-quality genomes of the isolate collection using the *de novo* SPAdes assembler (v3.11.1) (7). Re-constructed genomes were then subjected to *in silico* MLST profiling using the mlst tool (v2.23.0) (https://github.com/tseemann/mlst) with both the ‘Oxford’ and ‘Pasteur’ schemas for *A. baumannii*. Novel allele profiles and variants were deposited at PubMLST (https://pubmlst.org/). Additional AMR gene profiling was conducted through the AMRFinderPlus pipeline (v3.11.26) (https://github.com/ncbi/amr), utilizing the NCBI Antimicrobial Resistance Library for AMR (dated 2023-07-05) with default values of 80% for identity and coverage, respectively. Next, IS element characterization was performed via the ABRicate software (v1.0.1) (https://github.com/tseemann/abricate), utilizing a custom database based on the ISfinder collection (106) (dated 2023-07-05) with 90% identity and coverage. Finally, novel *bla*_ADC_ variants and the continuity of the *com*M gene were further investigated through a BLAST-based custom Python script. A total of 32 novel *bla*_ADC_ variants were identified and subsequently submitted to NCBI.

### Phylogenetic characterization

Open reading frames (ORFs) predicted by Prokka (v1.13) (104) were subsequently used as input for Roary (v3.12.0) (89) in order to conduct pan-genome analyses. Computed core genes were subsequently extracted, aligned and concatenated using default settings. The resulting alignment was then utilized to calculate a maximum likelihood-based phylogeny with RAxML (v.8.2.10) (110) using 100 bootstraps under the assumption of the GTR-gamma DNA substitution model. ClonalFrameML (v1.11) (19) was then used to correct for recombination events and phylogenetic groups were identified through Bayesian Analysis of Population Structure (BAPS). Here, we utilized BAPS with hierarchical clustering as implemented in the R package RhierBAPS (v1.0.1) (119). Grouping of the accessory genome was further assessed via t-distributed stochastic neighbor embedding (t-SNE), in order to cluster the data through a range of values for perplexity (p=5,10,20,50), the results of which were visualized using the microreact platform (4). SNP distances between individual samples were computed through the snippy pipeline (v4.6.0) (https://github.com/tseemann/snippy).

### Molecular clock analysis

In order to assess the molecular clock of *A. baumannii,* a xml file was configured using BEAUTi and subsequently utilized for BEAST (v2.5.0) (13) analysis. BEAST was run on the OXA−126 group, containing detailed sampling dates of 30 isolates. The run was allowed to continue for 50 million iterations, sampling from the posterior every 1000st iteration. We inferred the temporal phylogeny for OXA−126 under a strict clock setting (i.e. normally distributed rate variation over the tree). The strict clock was set to utilize a log-normal prior on the clock rate, with a mean of −5 and sd of 1.25 (43).

### Isolation of fungi and interaction studies with *A. baumannii*

Fungal isolates were obtained from environmental samples previously tested positive for *A. baumannii* using the following protocol: Samples (e.g. 1 g of soil) were resuspended in 10 ml of minimal salt medium supplemented with 0.2% acetate (see above) and 100 µl suspension before being spread on Kimmig fungal agar ((54), Becton Dickinson, Heidelberg, Germany) supplemented with 80 mg/l chloramphenicol (“Kimmig agar”). After 2−3 days of incubation at 27°C, fungal colonies were selected and spread on Kimmig agar several times until considered pure cultures after macroscopic and microscopic inspection. The absence of bacterial contamination was verified via colony PCR using global 16S rRNA gene primers (125). Provisional species determination of fungal isolates was performed by sequencing of the rRNA genes and ITS (internal transcribed spacer) (8, 118). *Aspergillus* spores were harvested after growth of fungi on Kimmig agar for 3 days at 34°C. To this end, the plate was flooded with 5 ml of sterile phosphate-buffered saline (PBS) supplemented with 0.1% Triton X−100, followed by soft panning. Subsequently, the suspension was removed with a pipette while avoiding contact with the fungal colonies, aliquoted and frozen at −80°C until further use. For seeding into 24-well plates, frozen spore suspensions were thawed, diluted by a factor of 50 into LB medium and 300 µl deposited into each well. The *A. baumannii* strains were cultured in LB medium overnight at 37°C, diluted 1:50 into fresh medium and further cultured until an optical density (OD) _600 nm_ of 1 was reached. For the spore adhesion assay, the bacteria were diluted 1:200 into LB medium and 300 µl were added to the spore seeding of a well. After 2−4 hours, the medium was removed from the wells and replaced twice with fresh medium to filter any non-adherent bacteria before inverted microscopy was performed. For the spore germination inhibition assay, bacterial suspensions with an OD _600 nm_ of 1 generated as above were diluted 1:2000 into LB medium, of which 300 µl were added to the spore seeding of a well. Microscopic evaluation of spore germination inhibition was conducted after 5−7 hours. For scanning electron microscopy (SEM), Thermanox^TM^ coverslips (ThermoFisher Scientific) were placed in 6-well plates, the incubation volume was reduced to 300 µl with seeding as above and fixed after 2 hours in a solution of 2.5% glutaraldehyde, 1.0% paraformaldehyde in 50 mM HEPES buffer for 24 hrs. All samples were then washed in 50 mM HEPES, dehydrated in 30, 50, 70, 90, 95, 100, 100% ethanol, critical point dried, mounted on aluminum stubs, sputter coated with a 10 nm layer of gold-palladium and finally examined in the SEM (ZEISS 1530 Gemini, Carl Zeiss Microscopy GmbH, Germany) operating at 3 kV using the in-lens electron detector.

### Geographic information analysis

All spatial data were analyzed using geographic information system (GIS) tools (Quantum GIS Software, 2010). Feeding grounds for all white stork nests were set as an area around the nest with a 3.5 and 4 km radius, respectively (11). Habitat structures of all feeding grounds were determined using CORINE Land Cover 2012 types, including arable land, pastures, heterogeneous agricultural areas, forests, total area of water reservoirs and total length of rivers. All results were combined for statistical analysis (Suppl. Material S1)

### Legal permissions

Collection of earthworms and insects was granted by the Landesamt für Umweltschutz, Sachsen-Anhalt, Halle/Saale, Germany (RL-0489-V and RL-0497-V). Collection of stork samples was granted by The Local Ethics Committee and The General Directorate for Environmental Protection in Poland: 21 21/2028, 42/2018, 43/2010, 44/2015, 47/2017, 028/2019/P1, 348/2016, DOP-OZGIZ.6401.03.101.2012.km.2, WBA/11/Z/10, WPN.6401.109.2017.AP, WPN.6401.211.2017.MK, DZP-WG.6401.75.2022.WW, DL-III.6713.11.2018.ABR, WPN-II.6401.167.2015.AS.2, DZP-WG.6401.03.98.2016.km, DLP-VIII−6713-21/29762/14/RN. In accordance with the convention on biological diversity (“Nagoya protocol”) a certificate of compliance was issued to collect samples from Spanish storks (ABSCH-IRCC-ES-246587−1; Dirección General de Biodiversidad y Calidad Ambiental del Ministerio para la Transición Ecológica, Madrid, Spain). Trapping small mammals in Thuringia and trapping Norway rats in North Rhine-Westphalia was covered by permits from the districts’ and State’s authorities, respectively (Thuringia: Landratsamt Unstrut-Hainich-Kreis file number 31109−16−301, Landkreis Eichsfeld file number 70.2−6-85/KleinSäuger/ausn.-01, Landratsamt Kyffhäuserkreis file number lll.3.3 364.53.1/2016−11-01; North Rhine-Westphalia: LANUV file number 84-02.04.2015.A279).

### Collection of rodents, rats and shrews

Cat-captured rodents and shrews were wrapped into aluminum foil or transferred into 50 ml tubes by the instructed collectors, stored at 4°C and transported to the lab within 24 h. Tracheal and rectal swab samples were taken under the safety cabinet while wearing gloves.

## Supporting information

Supplementary Excel sheets

## Acknowledgements

We would like to thank our colleagues at the central DNA sequencing lab of the Robert Koch Institute for excellent technical assistance. G.W. & P.G.H. acknowledge financial support from the Deutsche Forschungsgemeinschaft (DFG-FOR 2251). A. Ł. acknowledges financial support from a traineeship within the ERASMUS+ Programme (agreement no. 20/ SMP/2015/16). The collection of rats was funded by Deutsches Zentrum für Infektionsforschung (DZIF), thematic translational Uni (TTU) “Emerging Infections” to RGU; the collection of other rodents was commissioned and funded by the Federal Environment Agency (UBA) within the Environment Research Plan of the German Federal Ministry for the Environment, Nature Conservation, Building and Nuclear Safety (JJ, grant numbers 3716484310 and 3714674070). This publication made use of the *Acinetobacter baumannii* MLST website (http://pubmlst.org/abaumannii/) hosted at the University of Oxford (50). The development of this site has been funded by the Wellcome Trust. We would also like to thank numerous colleagues, friends and extramural partners supporting collection of sample material: Wolfgang Witte, Steffi Walter, Christina Lang, Christa Cuny, Ulrich Brandt, Alexander Riess, Magdalena Heindorf, Christian Schumacher, Andreas Tille, Emil Friedrich, Christian Imholt, Mechthild Budde, Engelbert Kampling, Angela Langbein-Lorf, Denise Kreuzmann, Christian Fritzsch, Reinhard Kappe, Bernd Nicolai. We are indepted to Mechthild and Christoph Kaatz (Storchenhof Loburg) for various support in obtaining stork samples, and to Volker Rickerts for support with fungal taxonomy. We would like to apologize to all those whose work could not be cited due to the extensive linkage of issues to the existing literature. We will try to acknowledge this in a review article.

