## Supplementary Figures for "On the ecology of *Acinetobacter baumannii* – jet stream rider and opportunist by nature"

**Suppl. Fig. S1** **Sampling of pellets, feathers and egg shells from white storks in Loburg (Germany) in 2015 and 2016**

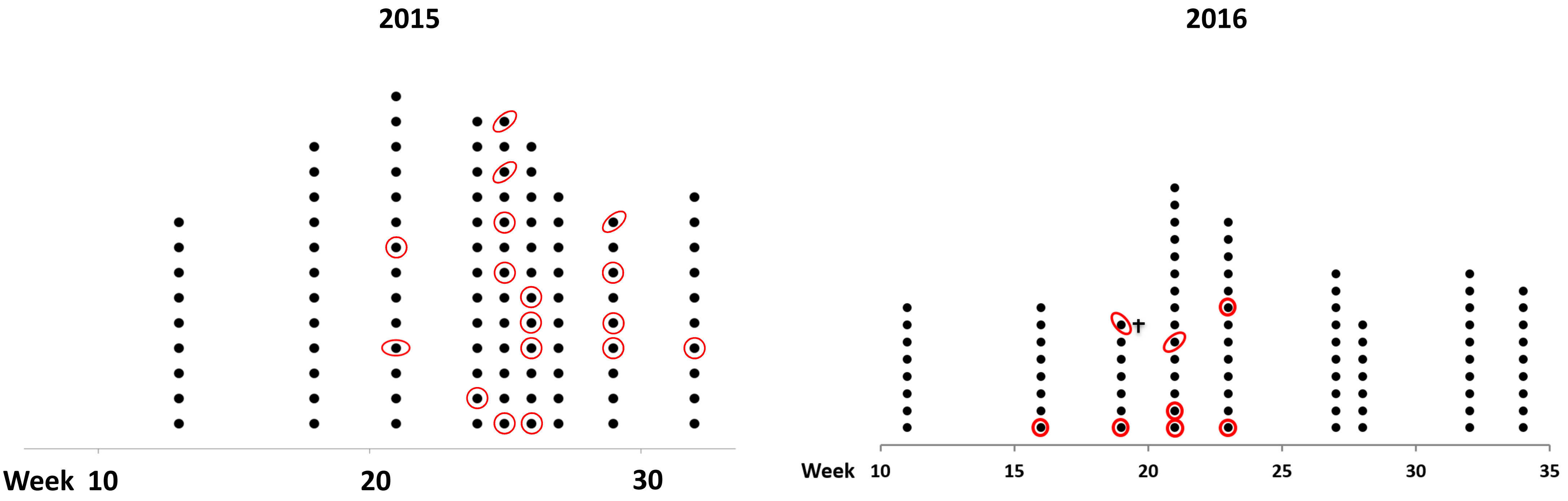


**Pellets of white storks were collected in 2015 and 2016 below nests in Loburg/Germany. Sporadically found feathers and egg shells were also sampled. Pellets from which *A. baumannii* was isolated are indicated by red circles. Red ellipses indicate egg shells and feathers tested positive, a single chick found dead was also tested positive and indicated with †.**

**Suppl. Fig. S2** 

### Pellets from white storks collected 2015 in Spain

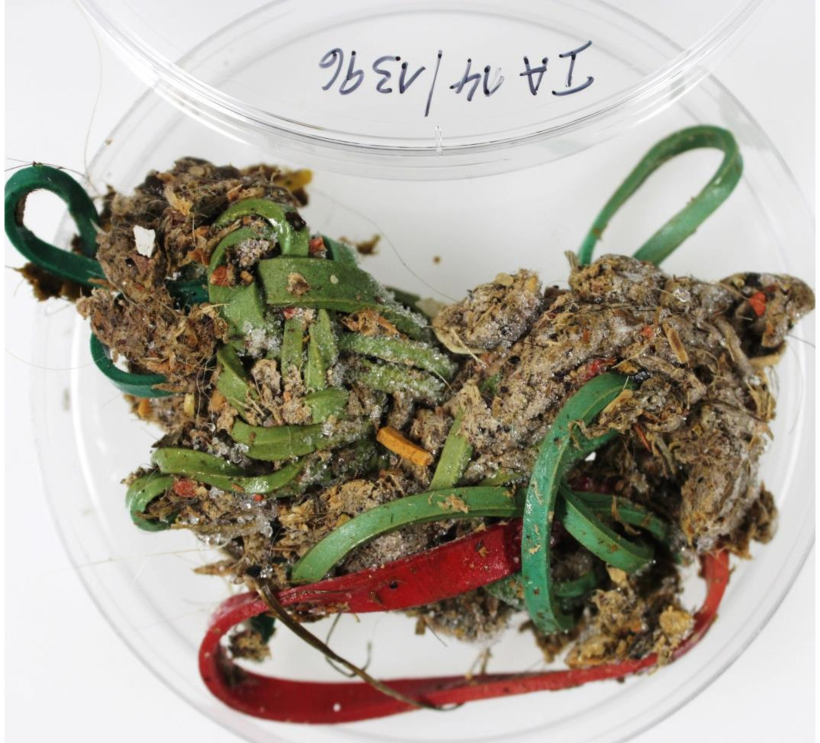

White stork‘s pellet from garbage dump

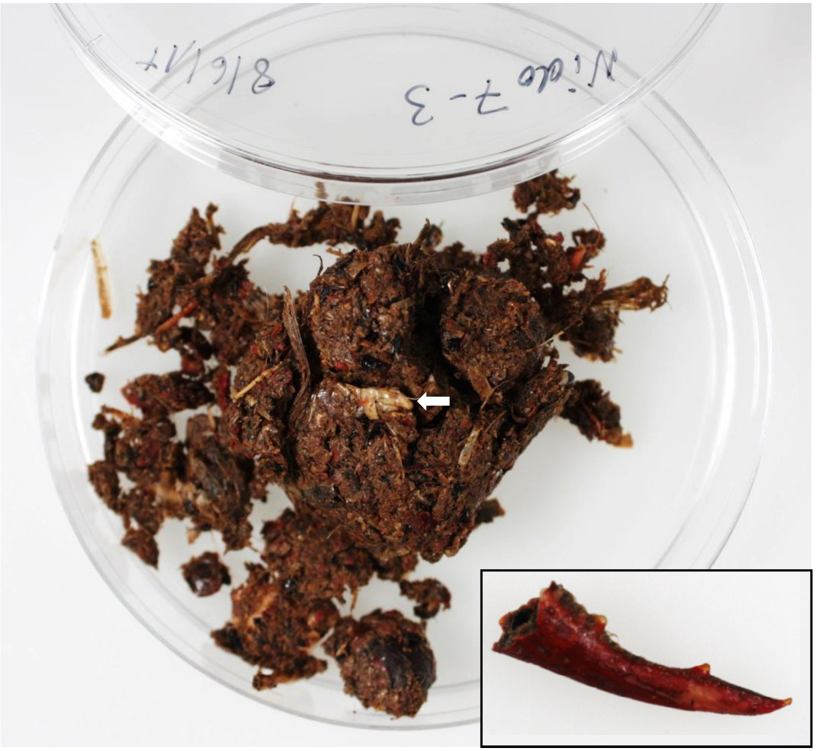

Pellet of white stork feeding in a natural environment.
Major identifiable feed: crabs (inset) and grasshoppers (arrow)

**Suppl. Fig. S3** **Outer and inner surface of a stork pellet copied on CHROMagar Acinetobacter**

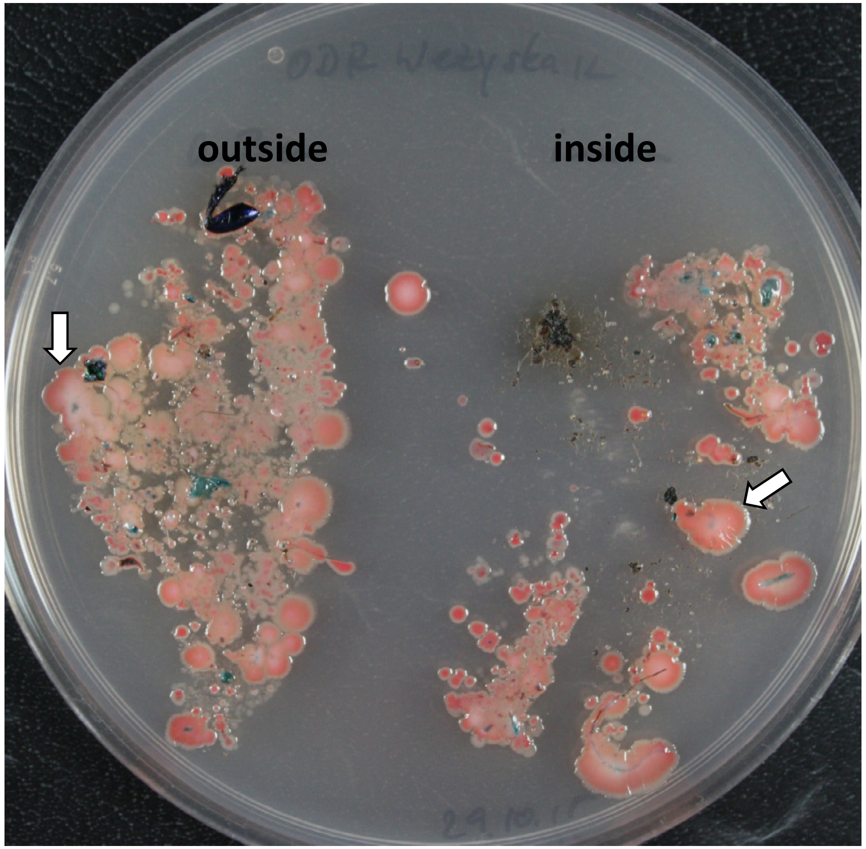


**Colonies with typical appearance of *A. baumannii* (see arrows) were confirmed by PCR.**

**Suppl. Fig. S4**

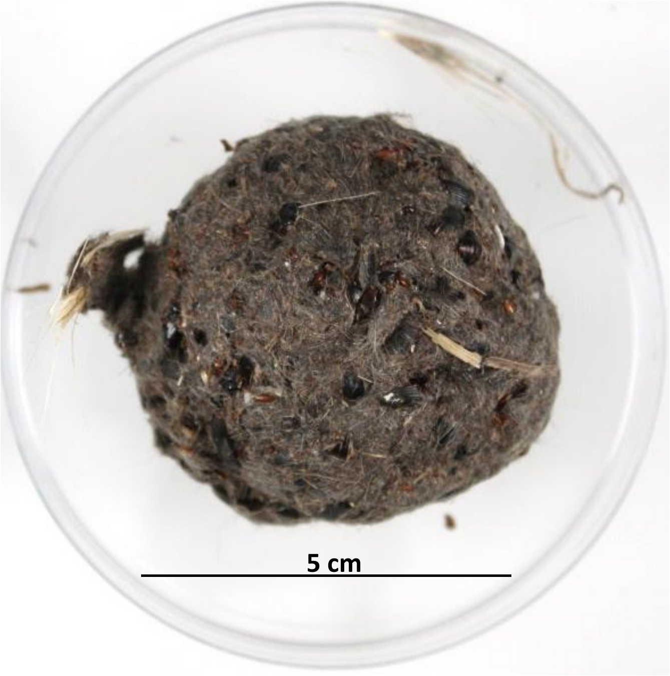


**Pellet of a white stork**

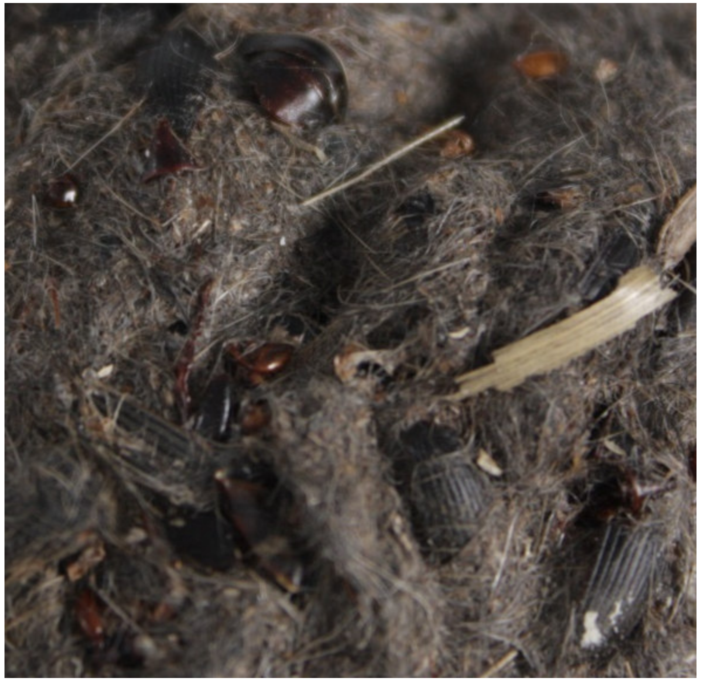

**Detail on a pellet of a white stork revealing animal hair and beetles' elytrae**

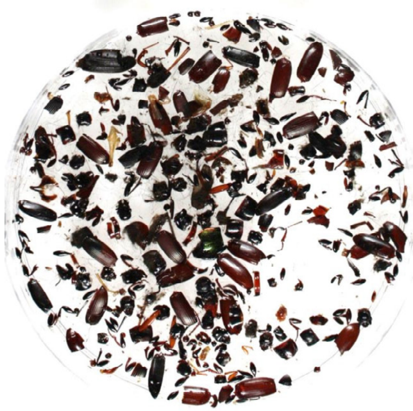

**Remains of arthropods from a stork pellet**

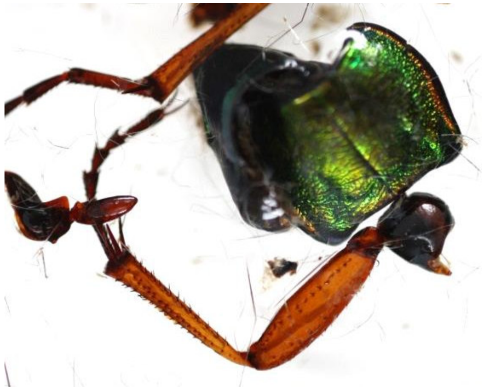

**Remains of *Carabus auratus* from a stork pellet**

**Suppl. Fig. S5** **Arthropod content of white stork pellets does not differ between pellets tested positive and negative for *A. baumannii***

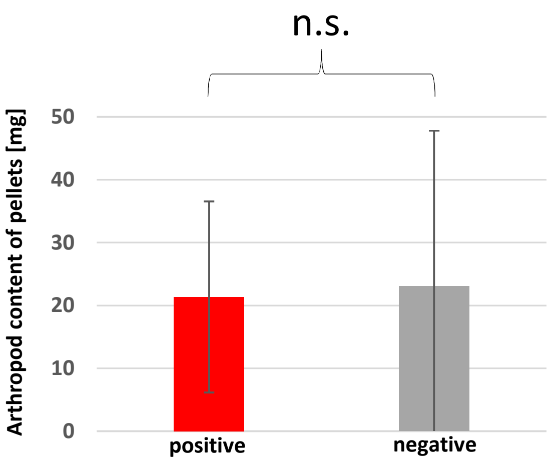


**Ten pellets each, tested positive and negative for *A. baumannii*, were dried until constant weight, and 1 g each dissected for arthropod content, which was weighed.**

**Suppl. Fig. S6** **Dominant arthropod content of white stork pellets collected in Loburg (Germany)**

***Zabrus tenebrioides***

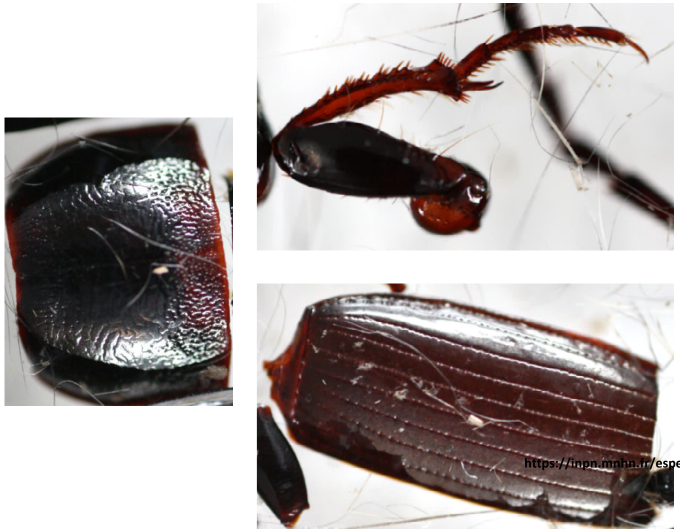

https://inpn.mnhn.fr/espece/cd_nom/222558

***Pterostichus melanarius***

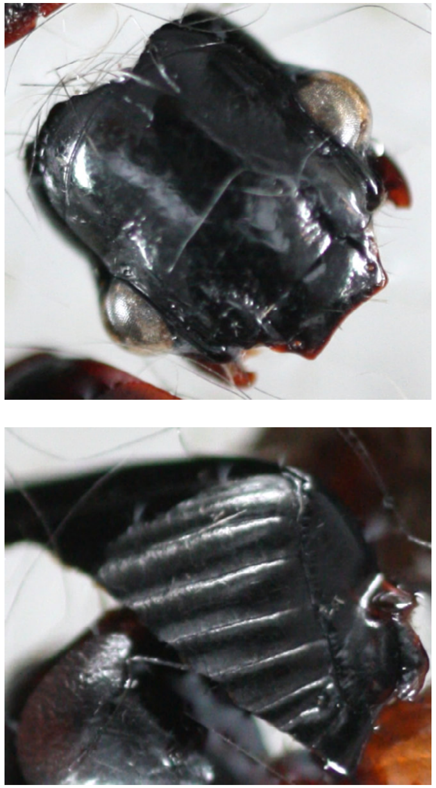

**Suppl. Fig. S7** **Cat-captured rodents and shrews tested for *A. baumannii* in 2014 and 2015 in Germany**

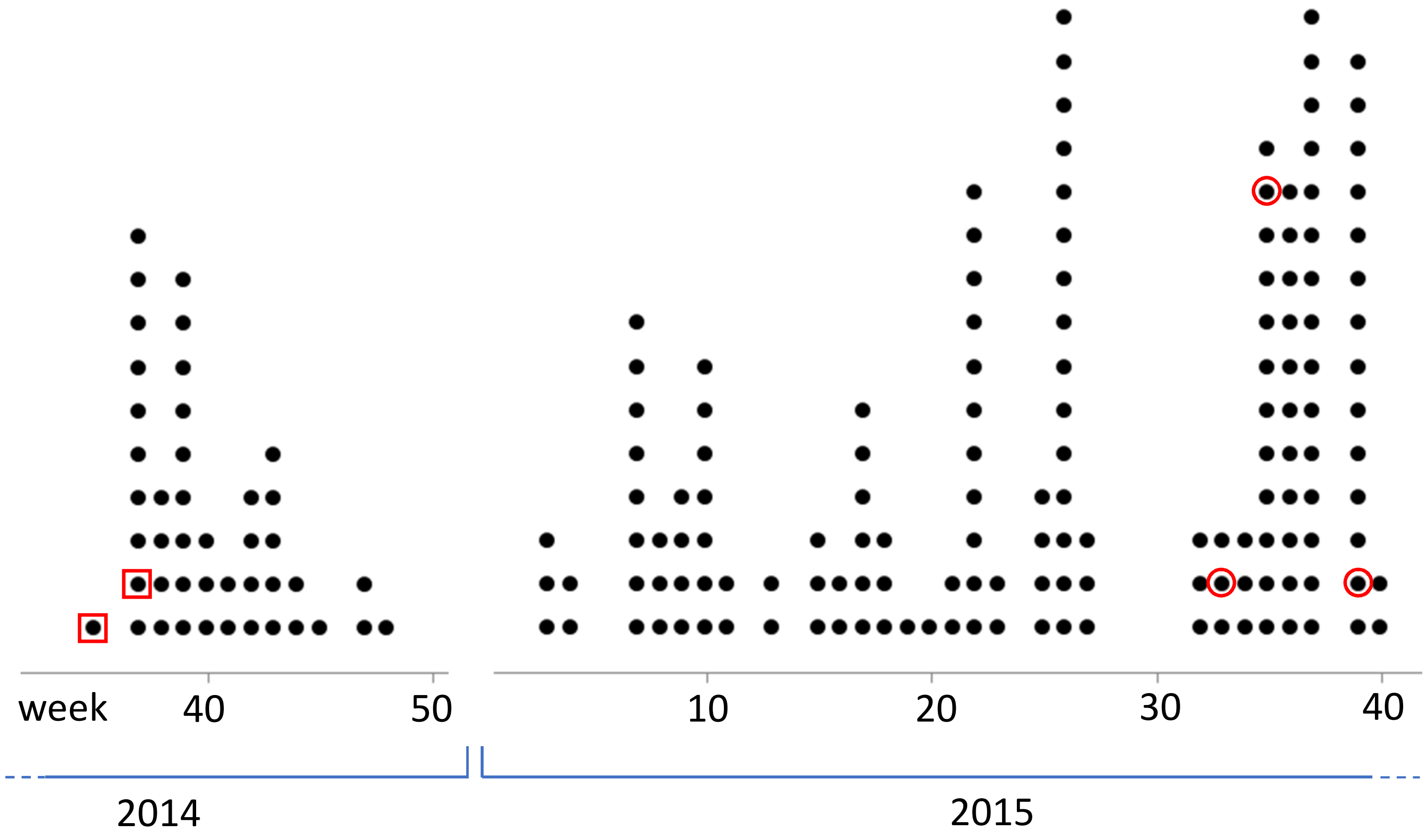


**Cat-captured rodents and shrews analysed in the years 2014 and 2015. Shrew from which *A. baumannii* was isolated are indicated by red squares, rodent tested positive indicated by a red circle.**

### Suppl. Fig. S8

### Grey heron pellets and egg shells tested for *A. baumannii*

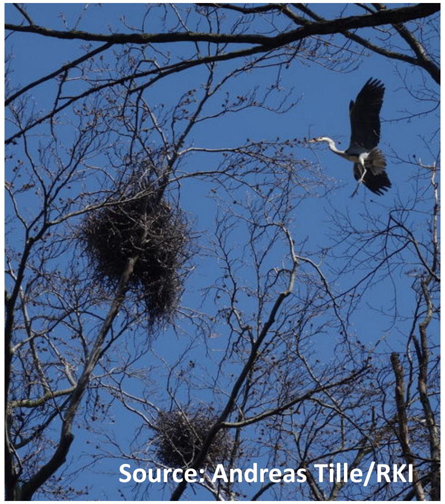


Grey heron approaching a nest

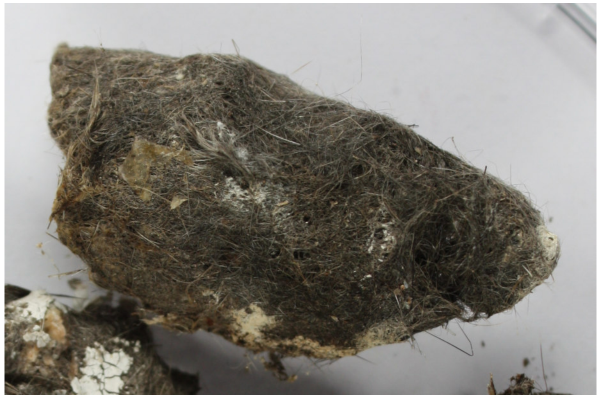

Pellet from grey heron

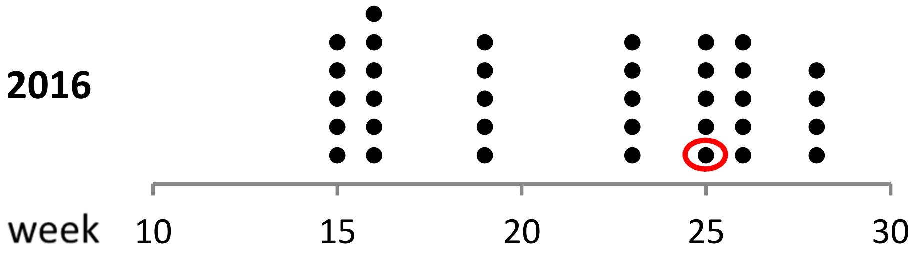


Pellets and egg shells from grey heron (*Ardea cinerea*) collected in Wernigerode (Germany) in 2016. Each black circle represents a single pellet or egg shell. An egg shell from which *A. baumannii* was isolated in June is indicated by a red ellipse.

**Suppl. Fig. S9**

### Kestrel pellets tested for *A. baumannii*

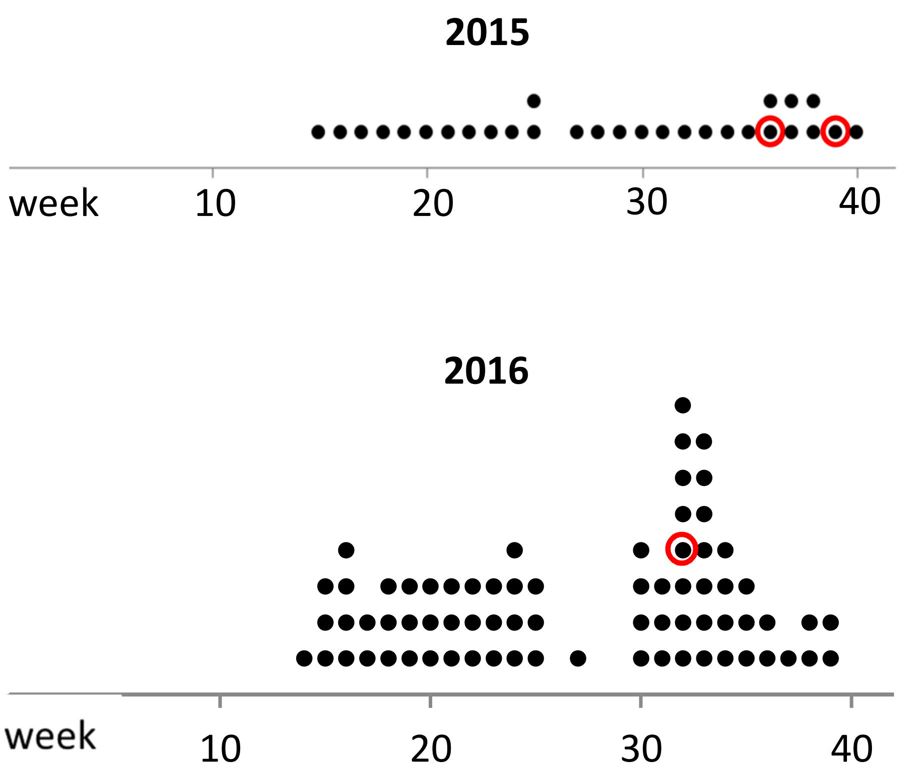


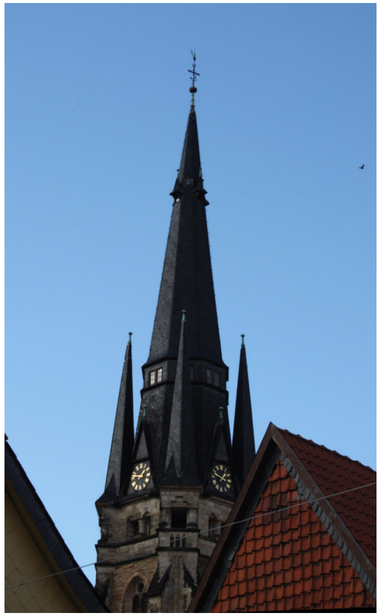

**Steeple of „Liebfrauen“ in Wernigerode, where pellets have been collected around the tower**

**Pellets from kestrel (*Falco tinnunculus*) collected in Wernigerode (Germany) in 2015 and 2016. Each black circle represents a single pellet. Pellets from which *A. baumannii* was isolated are encircled red.**

Suppl. Fig. S10

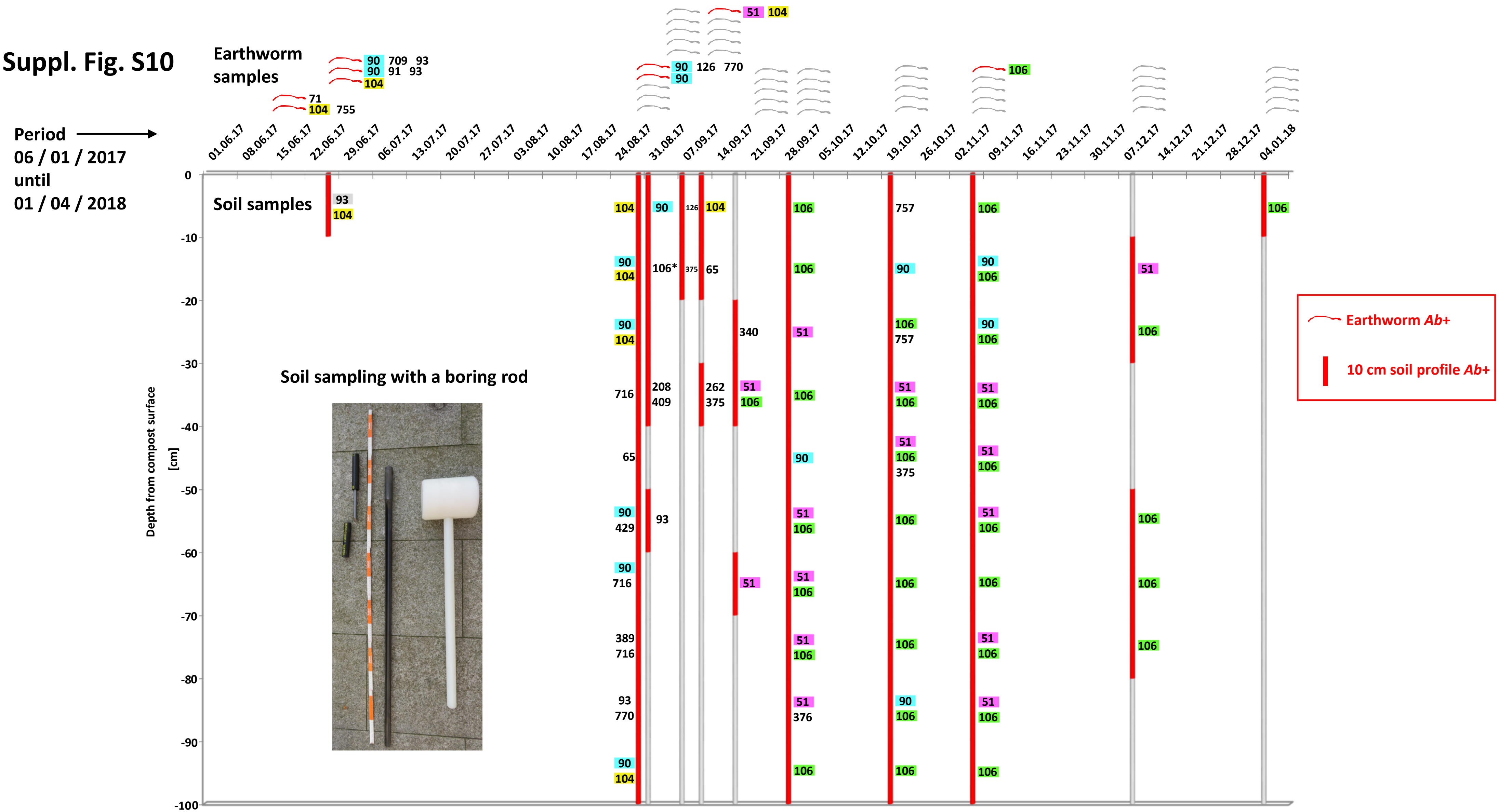


Sampling of compost soil and earthworms in the same compost in Wernigerode (Germany) over time. Numbers indicate OXA-types of *A. baumannii* isolates.

### Suppl. Fig. S11

### Sampling the Holtemme river bank

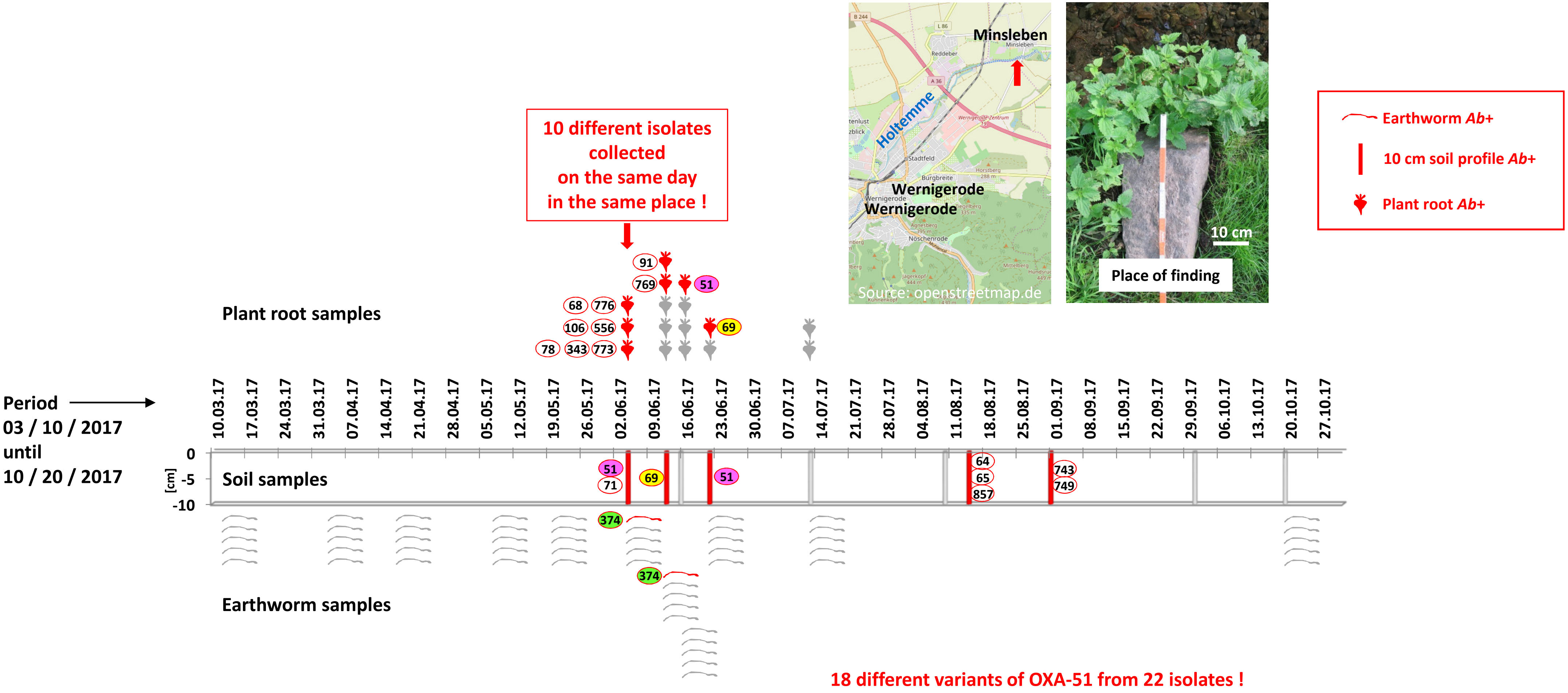


Sampling site at the river Holtemme near Minsleben/Wernigerode (Germany). Numbers indicate OXA-types of *A. baumannii* isolates.

**Suppl. Fig. S12** **Soil samplings in the forest south of Wernigerode/Germany (2016-2021)**

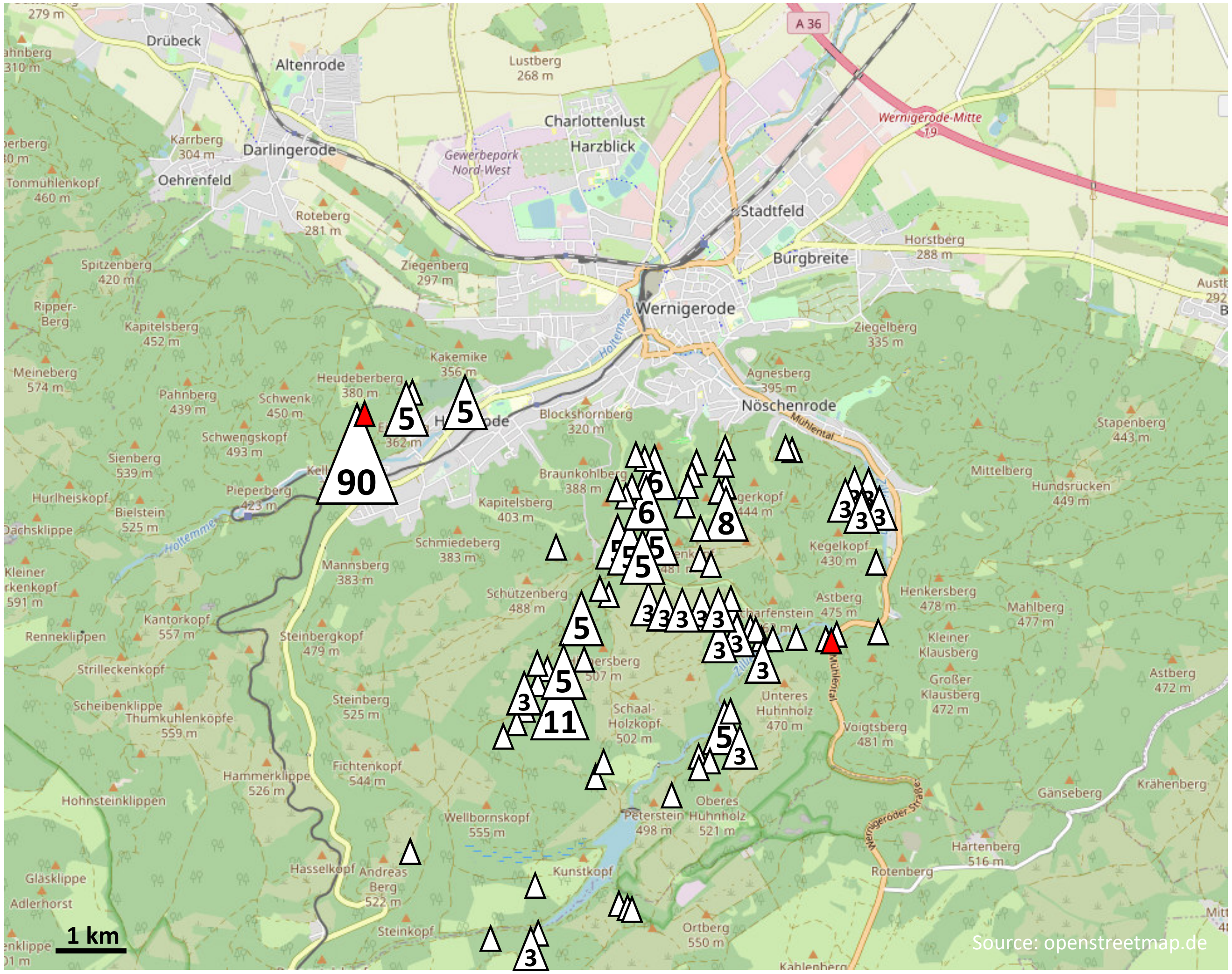


**Number of soil samples at a specific site indicated if >1; samples from which *A. baumannii* was isolated indicated in red; ∑=269 samples, 2 positiv (0.7%)**

**Suppl. Fig. S13** **Sampling at Benzingerode (08/30/2020 – 09/10/2021)**

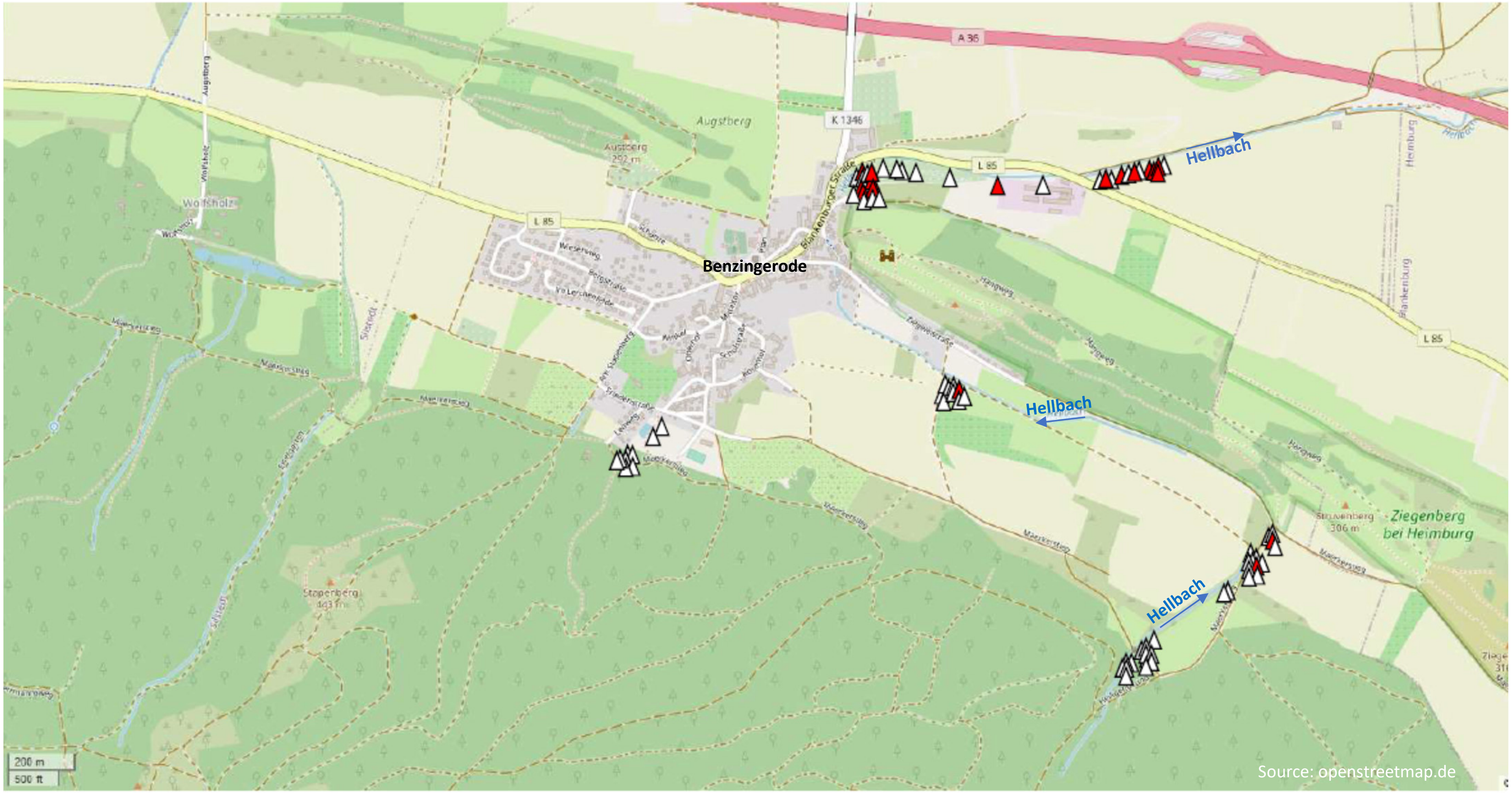


**No. of samples from the edge of the forest: 22 (all negative). Samples from which *A. baumannii* was isolated are indicated in red.**

**Suppl. Fig. S14** **Phylogenetic distribution of strains harbouring selected OXA-51-like variants**

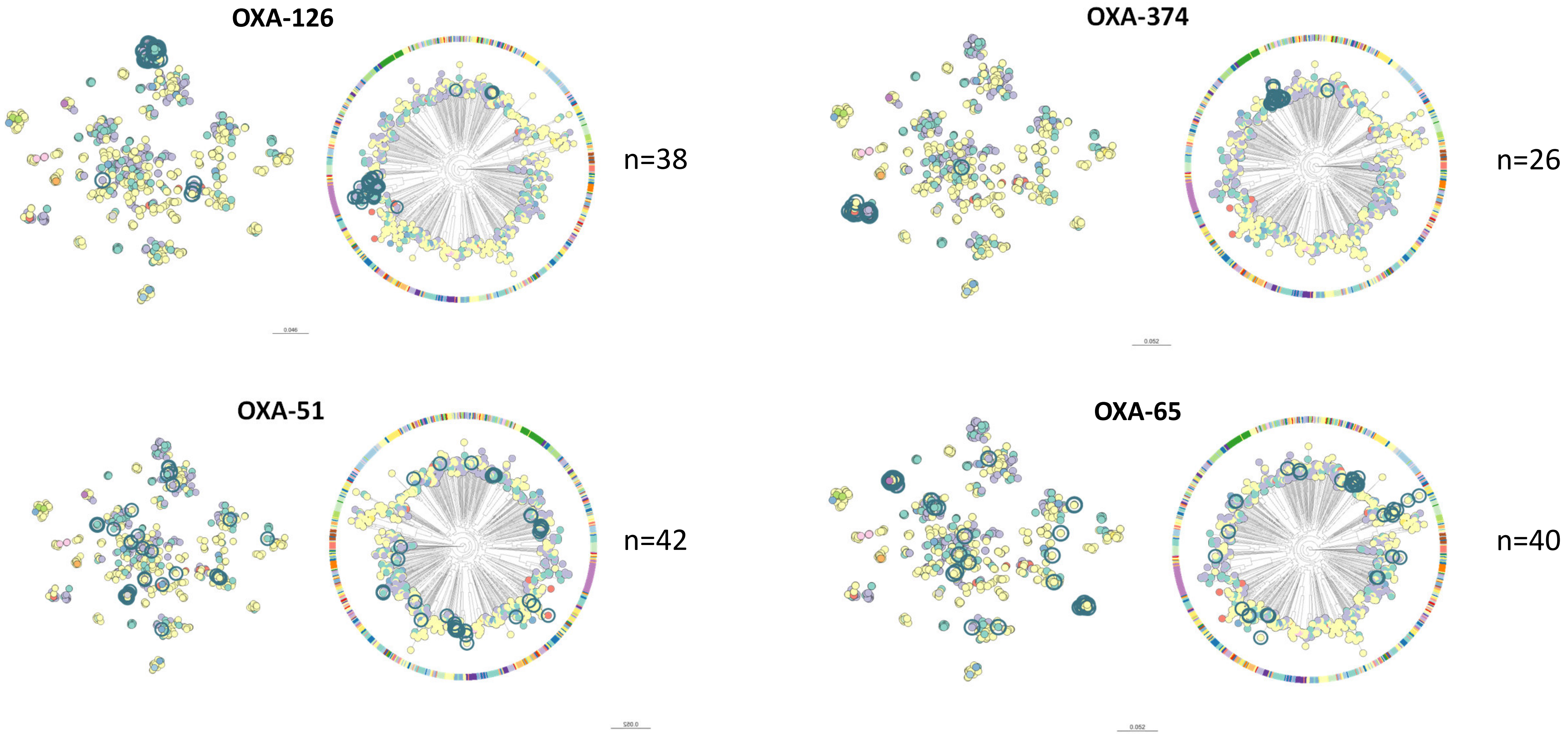


**See microreact project at [https://microreact.org/project/3ApuGKD61qPLT1ZNmoKcTb-acinetobacternobaps12dec23](https://microreact.org/project/3ApuGKD61qPLT1ZNmoKcTb-acinetobacternobaps12dec23) for further analysis and illustration of the dataset.**

**Suppl. Fig. S15** **Relevance of the One-Health concept using the example of international clone 8 (IC8)**

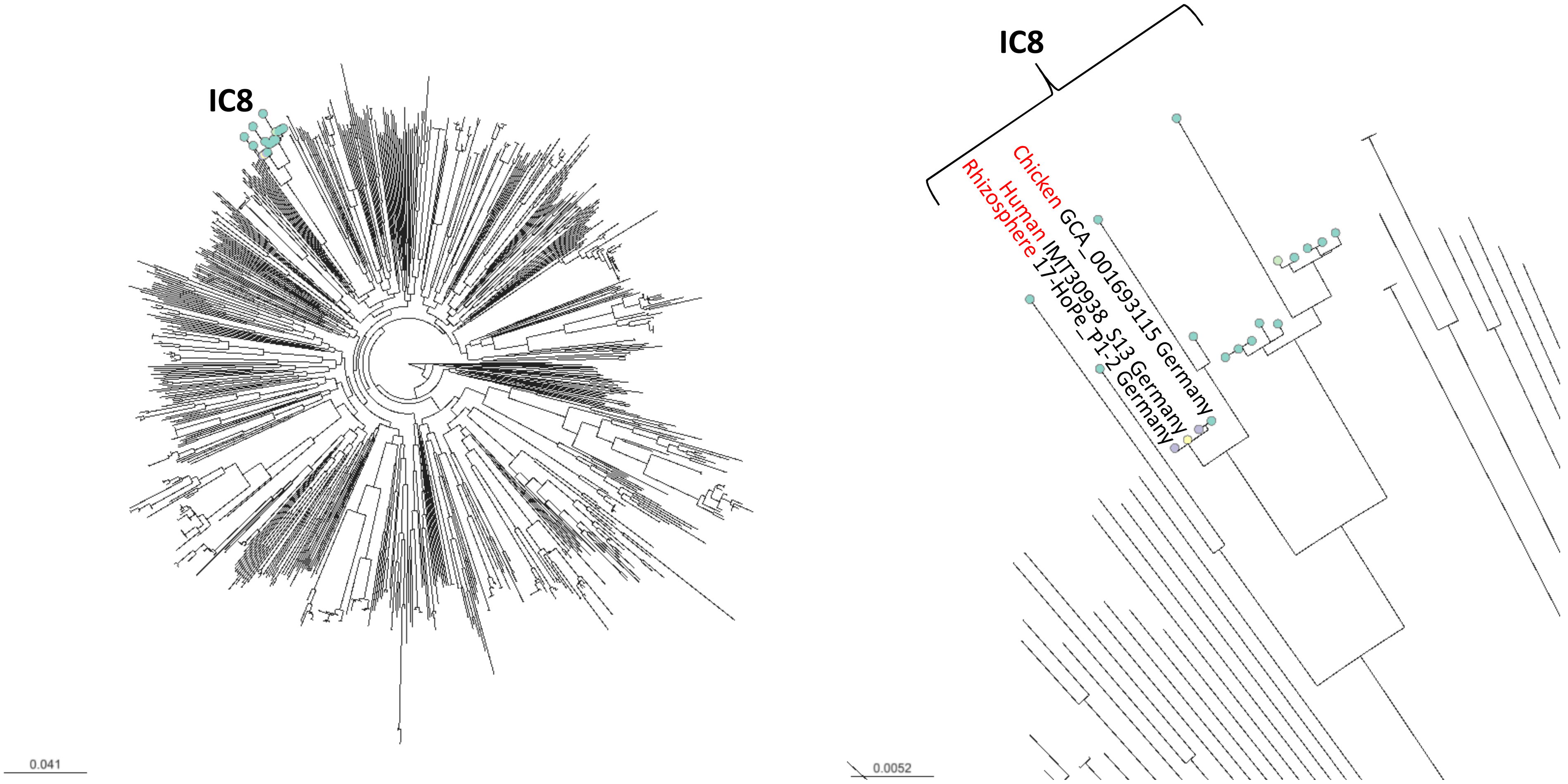

**Suppl. Fig. S16** **Penetrance of resistance genes among strains sequenced in this study**

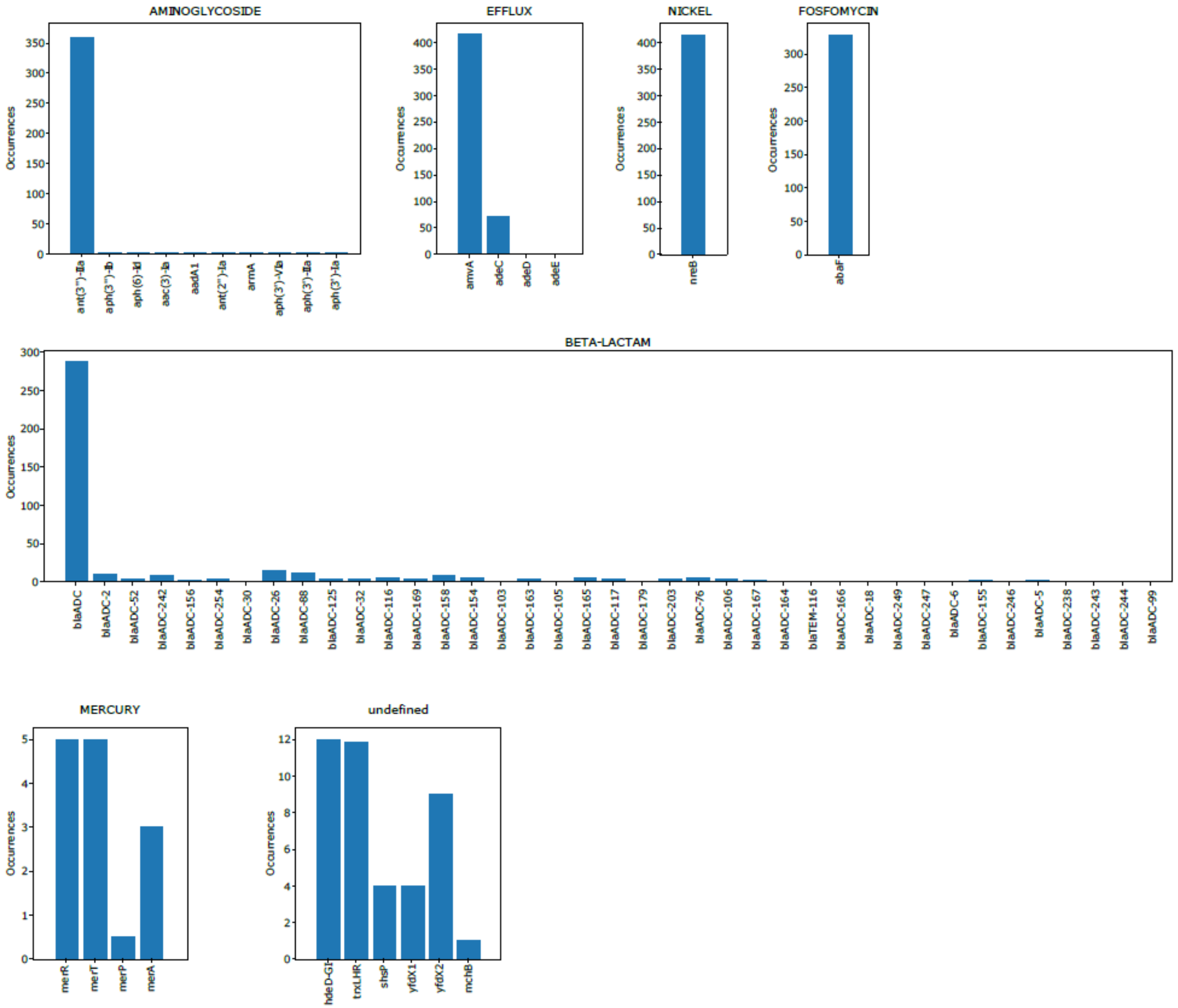


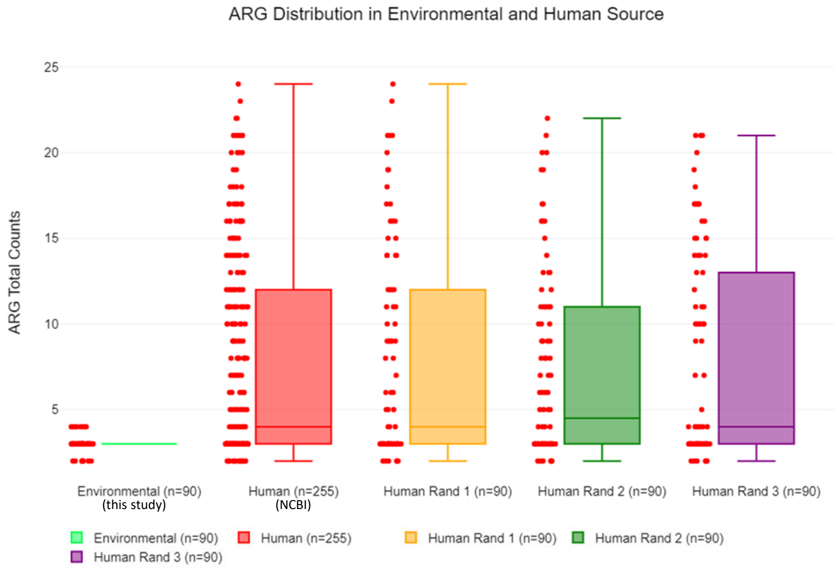


| Groups: | Environmental (n=90) | Human (n=255) | Human Rand 1 (n=90) | Human Rand 2 (n=90) | Human Rand 3 (n=90) |
|---|---|---|---|---|---|
| Sample size (n): | 90 | 255 | 90 | 90 | 90 |
| Minimum: | 2 | 2 | 2 | 2 | 2 |
| Q1: | 3 | 3 | 3 | 3 | 3 |
| Median: | 3 | 4 | 4 | 4.5 | 4 |
| Q3: | 3 | 12 | 12 | 11 | 13 |
| Maximum: | 4 | 24 | 24 | 22 | 21 |
| Mean (x̄): | 3.077778 | 7.670588 | 7.711111 | 7.188889 | 7.744444 |
| Skewness: | 0.0998321 | 0.949572 | 1.011713 | 1.117898 | 0.825504 |
| Skewness Shape: | | | | | |
| Excess kurtosis: | 0.672117 | -0.360615 | -0.225786 | 0.178957 | -0.791584 |
| Tails Shape: | Potentially | Potentially | Potentially | Potentially | Potentially |
| Outliers: | 4, 2, 2, 4, 4, 2, 4, 4, 4, 4, 4, 2, 4, 4, 4, 4, 4, 2, 4, 4, 4, 2, 2, 2, 2 | | | | |

### Suppl. Fig. S17 IS element content in human and environmental isolates

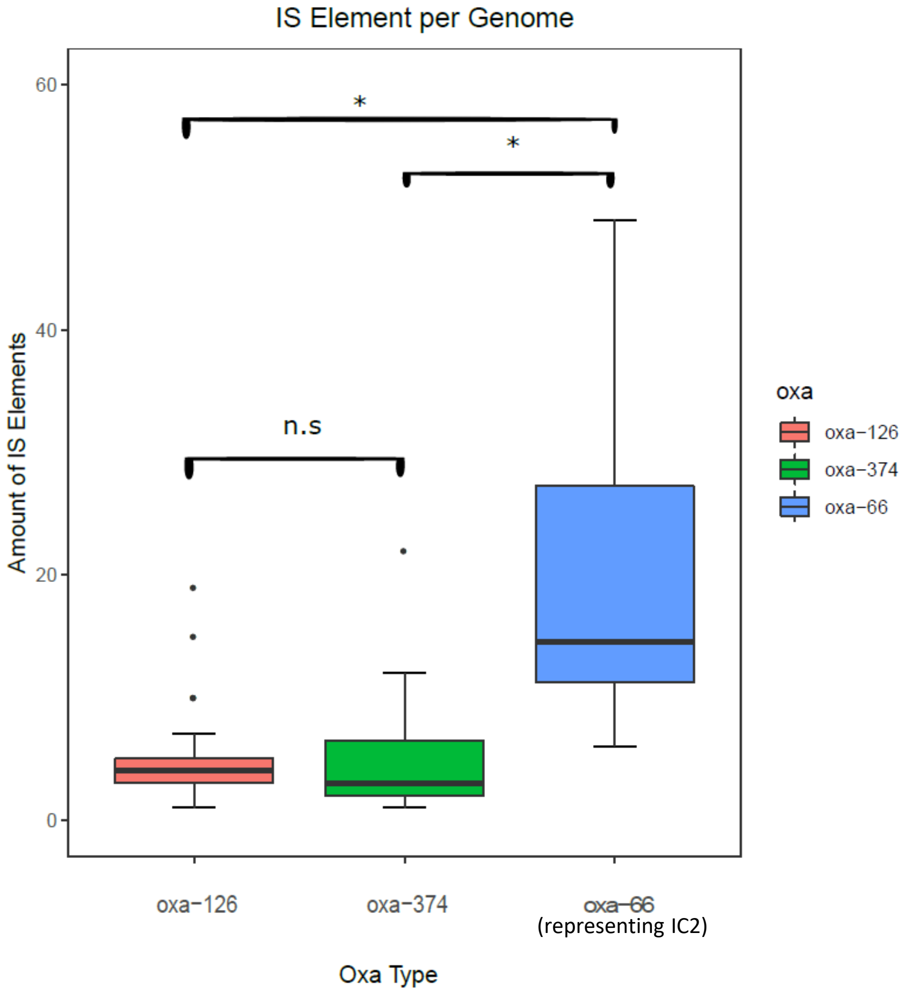


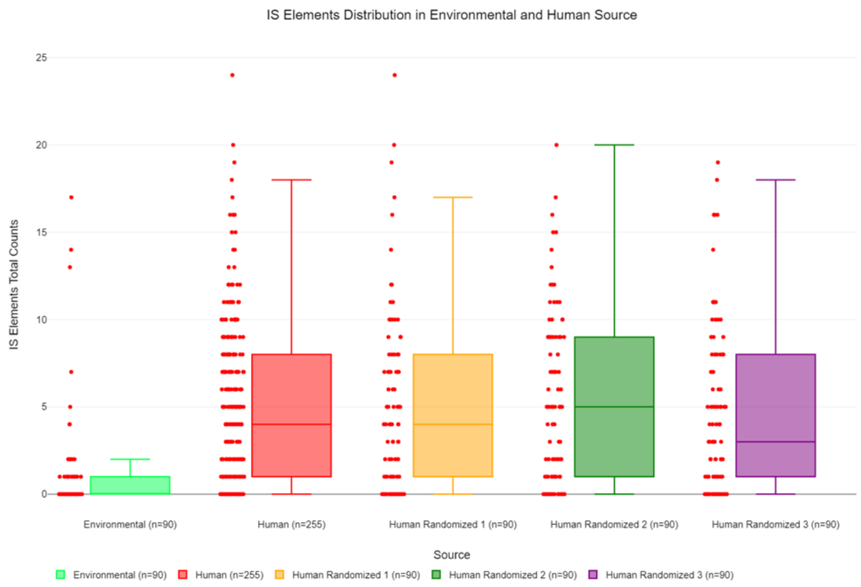


| Groups: | Environmental (n=90) | Human (n=255) | Human Randomized 1 (n=90) | Human Randomized 2 (n=90) | Human Randomized 3 (n=90) |
|---|---|---|---|---|---|
| Sample size (n): | 90 | 256 | 90 | 90 | 90 |
| Minimum: | 0 | 0 | 0 | 0 | 0 |
| Q1: | 0 | 1 | 1 | 1 | 1 |
| Median: | 0 | 4 | 4 | 5 | 3 |
| Q3: | 1 | 8 | 8 | 9 | 8 |
| Maximum: | 17 | 24 | 24 | 20 | 19 |
| Mean ($\bar{x}$): | 1.044444 | 4.667969 | 4.933333 | 5.211111 | 4.444444 |
| Skewness: | 4.223213 | 1.087683 | 1.364896 | 0.78632 | 1.18683 |
| Skewness Shape: | | | | Asymmetrical, | Asymmetrical, |
| Excess kurtosis: | 18.901693 | 1.12327 | 2.087564 | -0.123233 | 0.863476 |
| Tails Shape: | Leptokurtic, | Leptokurtic, | Leptokurtic, | Potentially | Potentially |
| Outliers: | 7, 17, 4, 14, 13, 5, 4 | 19, 20, 24 | 19, 24, 20 | | 19 |

**Suppl. Fig. S18**

### Geographic information systems (GIS) based studies on the foraging grounds around nests of white stork: dependency of *A. baumannii* colonisation on land use?

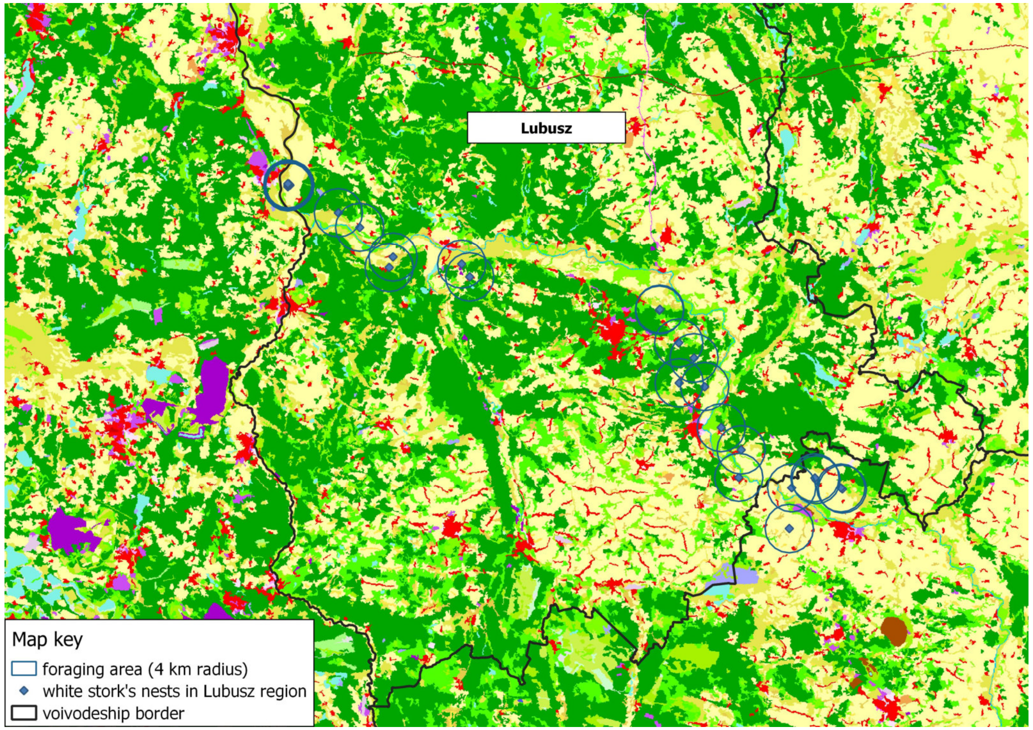


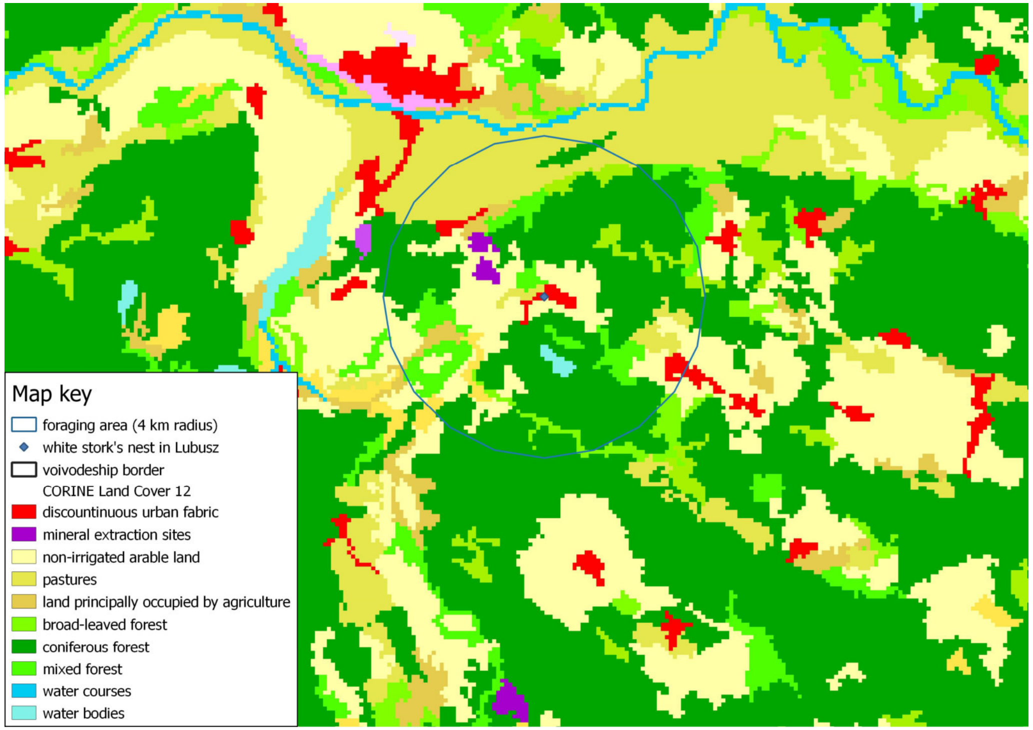


**Examplification of GIS based analyses of land use in the vicinity of white stork nests (foraging zone with a 4 km radius around nests). For further details see Suppl. Material S1.**

**Suppl. Fig. S19** **Probability of colonization of white stork nestlings with *A. baumannii* decreases with age of nestlings**

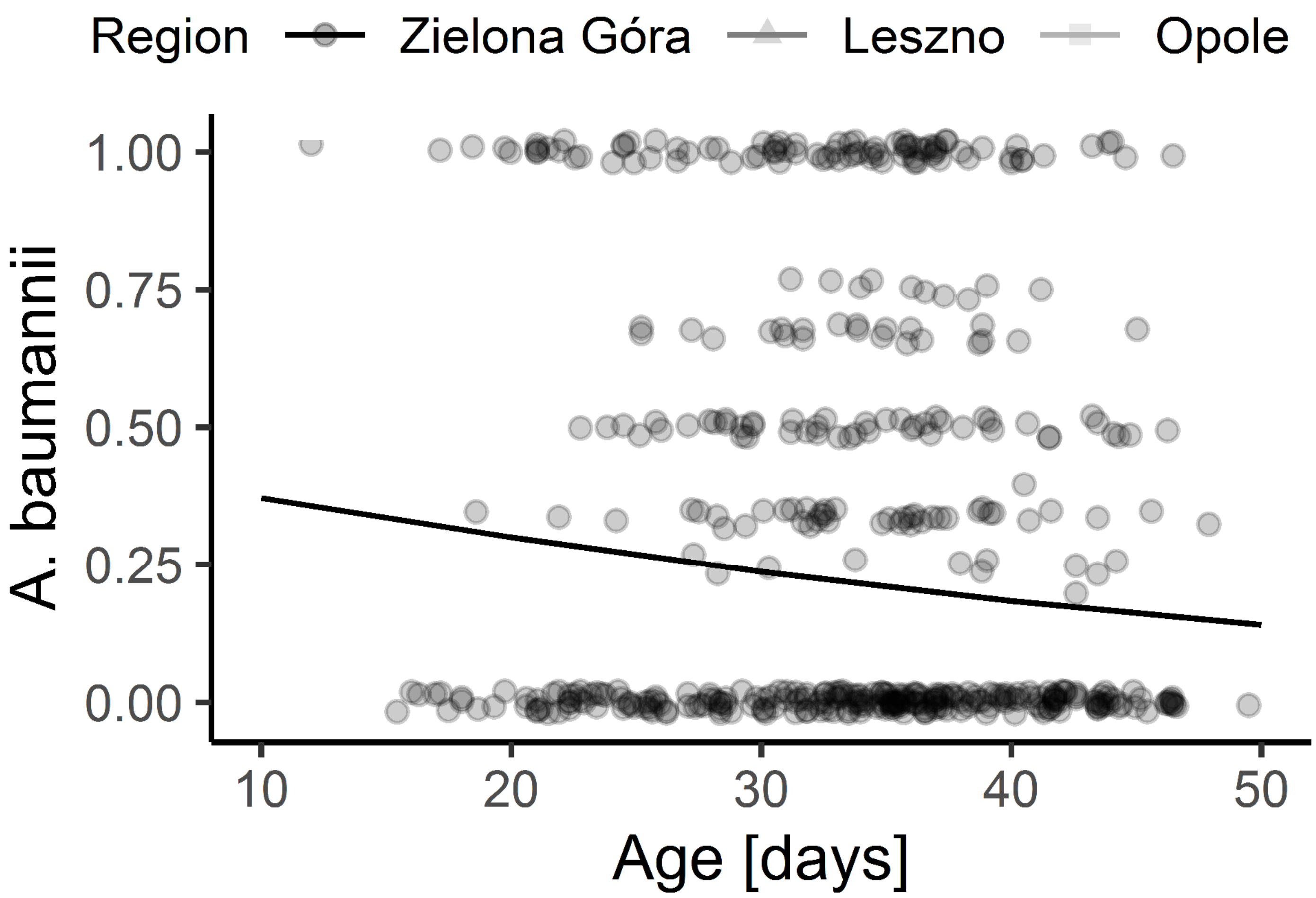


**For further details see Suppl. Material S1.**

### Suppl. Fig. S20 Extensive horizontal gene transfer among lineages co-existing in a natural habitat at river Holtemme

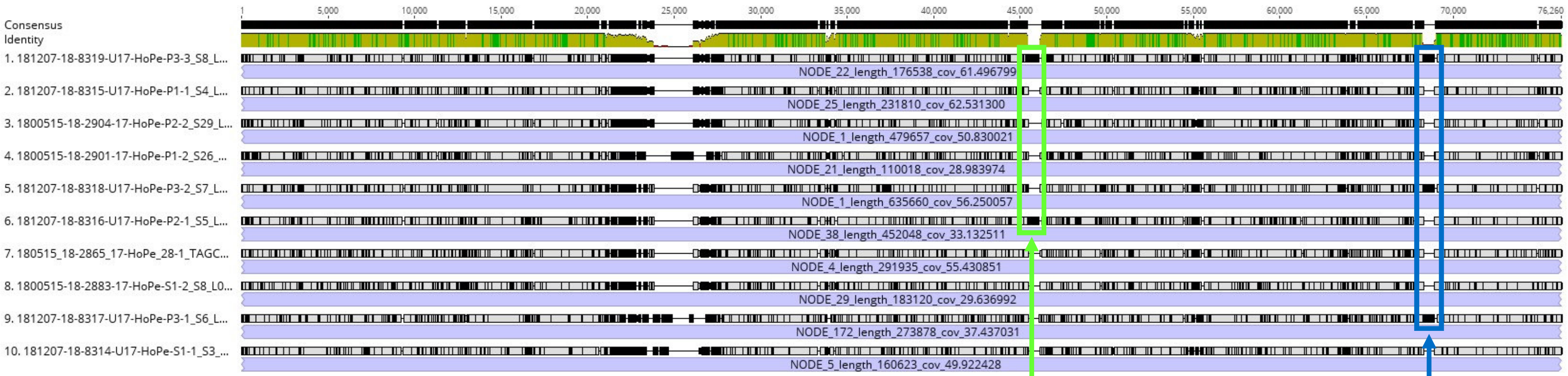


**SMI1/KNR4 family protein**

**Only 1 nucleotide BLAST hit in *Acinetobacter***

**N-Acetyltransferase**

**Only 22 nucleotide BLAST hits in *Acinetobacter***

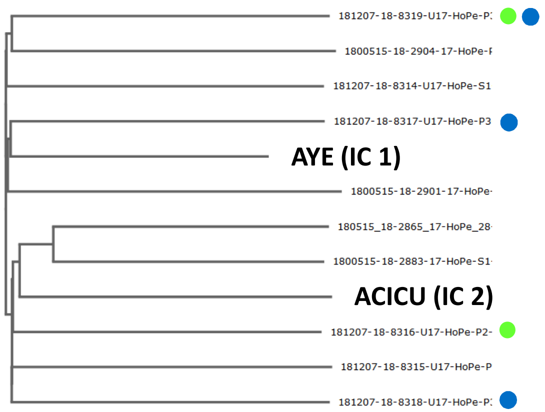


**Core genome-based phylogeny applying the BacWGSTdb web tool (Ruan et al., 2019)**

➡ **Horizontal gene transfer is a frequent event in the nature**
