## Supplementary text files for "On the ecology of *Acinetobacter baumannii* – jet stream rider and opportunist by nature"

**List of Supplementary Files**


Supplementary Figures

Suppl. Fig. S1: Seasonal studies on pellets of white storks in Germany

Suppl. Fig. S2: Content of white stork pellets from Spain

Suppl. Fig. S3: *A. baumannii* is not only on the surface but also inside of pellets from white stork

Suppl. Fig. S4: Content of white stork pellets from Germany

Suppl. Fig. S5: Arthropod content of white stork pellets does not correlate with *A. baumannii* isolation

Suppl. Fig. S6: Dominant arthropods found in pellets: ground beetles *Zabrus tenebrioides* and *Pterostichus melanarius*

Suppl. Fig. S7: Seasonal studies on *A. baumannii* isolation rates from cat-captured mice and shrews

Suppl. Fig. S8: Grey heron (*Ardea cinerea*) sampling

Suppl. Fig. S9: Kestrel (*Falco tinnunculus*) sampling

Suppl. Fig. S10: Sampling of garden compost

Suppl. Fig. S11: Sampling at the banks of river Holtemme near Minsleben

Suppl. Fig. S12: Sampling in the forest south of Wernigerode

Suppl. Fig. S13: Sampling at creek Hellbach near Benzingerode

Suppl. Fig. S14: Phylogenetic distribution of selected OXA-51-like variants

Suppl. Fig. S15: International clone 8 (IC8) includes environmental, animal and human isolates

Suppl. Fig. S16: Penetrance of resistance genes among strains sequenced in this study

Suppl. Fig. S17: IS element content in international clones and environmental isolates

Suppl. Fig. S18: Geographic information systems (GIS) based studies on the foraging grounds of white storks: dependency of *A. baumannii* colonisation on land use?

Suppl. Fig. S19: Probability of colonization of white stork nestlings with *A. baumannii* decreases with age of nestlings

Suppl. Fig. S20: Extensive horizontal gene transfer among lineages co-existing in a natural habitat

Supplementary Tables

Suppl. Table S1: Summary on samplings and isolates.

Suppl. Table S2: Blood parameters of white stork nestlings colonized or not with *A. baumannii.*

Suppl. Table S3: Novel OXA-51-like variants described in this study.

Suppl. Table S4: Metadata on strains selected for genome sequencing and strains selected from public databases (incl. MLST).

Suppl. Table S5: Distance matrix in terms of single nucleotide polymorphisms (SNPs) based on the genome alignment.

Suppl. Table S6: Remarkable relationships between strains described in this study and strains of public databases as well as among strains described in this study.

Suppl. Table S7: BEAST analysis on the OXA-126 clade.

Suppl. Table S8: Further evidence for the interrelatedness of the *A. baumannii* life cycle with the fungal world.

Supplementary Material

Suppl. Material S1: Supplementary information on GIS-based analyses.

### Suppl. Table S1:

### Summary on environmental/wildlife samplings and isolates

#### White stork nestlings

##### Poland (choana/trachea sampling) 2013-2018

| Region/voivodship | No. Choana samples | No. *Ab* positive samples | Percentage positive samples |
|---|---|---|---|
| Lubusz | 422 | 101 | 23.9 |
| Opole | 496 | 146 | 29.4 |
| Masovia | 80 | 12 | 15 |
| Greater Poland | 321 | 130 | 40.5 |
| **Poland total** | **1,319** | **389** | **29.5** |

##### Poland (rectal sampling) 2013-2018

| Region/voivodship | No. rectal samples | No. *Ab* positive samples | Percentage positive samples |
|---|---|---|---|
| Lubusz | 144 | 9 | 6.3 |
| Greater Poland | 183 | 31 | 16.9 |
| Opole | 340 | 18 | 5.3 |
| Masovia | 80 | 2 | 2.5 |
| **Poland total** | **747** | **60** | **8** |

##### Germany (choana/trachea) 2015

| Region | No. Choana samples | No. *Ab* positive samples | Percentage positive samples |
|---|---|---|---|
| **Germany total** | **29** | **10** | **34.5** |

##### Spain (choana/trachea) 2015 and 2019

| Region | No. Choana samples | No. *Ab* positive samples | Percentage positive samples |
|---|---|---|---|
| Spain 2015 | 57 | 0 | 0 |
| Spain 2019 | 64 | 5 | 7.8 |
| **Spain total** | **121** | **5** | **4.1** |

##### Spain (rectal sampling) 2019

| Region | No. rectal samples | No. *Ab* positive samples | Percentage positive samples |
|---|---|---|---|
| Spain 2019 | 64 | 0 | 0 |
| **Spain total** | **64** | **0** | **0** |

**Spain (pellets) 2015**

| **Region** | **No. pellet samples** | **No. *Ab* positive samples** | **Percentage positive samples** |
|---|---|---|---|
| Spain 2015 | 14 | 0 | 0 |
| **Spain total** | **14** | **0** | **0** |

**Spain (nest material) 2019**

| **Region** | **No. samples (swabs from nest material)** | **No. *Ab* positive samples** | **Percentage positive samples** |
|---|---|---|---|
| Spain 2019, natural nest material | 17 | 6 | 35.3 |
| Spain 2019, anthropogenic nest material (e.g. plastics) | 8 | 1 | 12.5 |
| **Spain total** | **25** | **7** | **28** |

**Spain (egg shells) 2019**

| **Region** | **No. egg shell samples** | **No. *Ab* positive samples** | **Percentage positive samples** |
|---|---|---|---|
| Spain 2019 | 60 | 0 | 0 |
| **Spain total** | **60** | **0** | **0** |

**Poland/**Greater Poland **(egg shells) 2015-2016**

| **Region** | **No. egg shell samples** | **No. *Ab* positive samples** | **Percentage positive samples** |
|---|---|---|---|
| Greater Poland 2015 | 42 | 18 | 42.9 |
| Greater Poland 2016 | 26 | 9 | 34.6 |
| **Poland total** | **68** | **27** | **39.7** |

#### Black stork nestlings

**Poland (rectal sampling) 2016-2020**

| **Region/voivodship** | **No. rectal samples** | **No. *Ab* positive samples** | **Percentage positive samples** |
|---|---|---|---|
| **Poland total** | **64** | **0** | **0** |

**Nest material (2020): 8 samples (all negative)**

#### Kestrel (*Falco tinnunculus*)

| Region/voivodship | No. pellet samples | No. *Ab* positive samples | Percentage positive samples |
|---|---|---|---|
| **Wernigerode (Germany)** | **101** | **3** | **3** |

#### Grey heron (*Ardea cinerea*)

| Region/voivodship | No. pellet, feather and egg shell samples | No. *Ab* positive samples | Percentage positive samples |
|---|---|---|---|
| **Wernigerode (Germany)** | **35** | **1 (egg shell)** | **3** |

#### Earthworms

| Region/voivodship | No. of earthworms | No. *Ab* positive earthworms | Percentage positive samples |
|---|---|---|---|
| **Saxony-Anhalt, mostly Wernigerode (Germany)** | **618** | **22** | **3.6** |

#### Soil & Rhizosphere (mostly Harz district, Saxony-Anhalt, Germany)

| Sample type | No. of samples | No. *Ab* positive samples | Percentage positive samples |
|---|---|---|---|
| Soil including compost 2017 | 181 | 88 | 48.6 |
| Rhizosphere 2017 | 77 | 15 | 19.5 |
| Soil including compost 2018 | 66 | 11 | 16.7 |
| Rhizosphere 2018 | 49 | 6 | 12.2 |
| Soil including compost 2019 | 179 | 39 | 21.8 |
| Rhizosphere 2019 | 62 | 17 | 27.4 |
| Soil including compost 2020 | 189 | 30 | 15.9 |
| Rhizosphere 2020 | 57 | 4 | 7 |
| Soil including compost 2021 | 155 | 28 | 18 |
| Rhizosphere 2021 | 21 | 6 | 28.6 |
| **Soil and rhizosphere total 2017-2021** | **1,036** | **244** | **23,6** |

Of these 1,036 soil samples: from compost and from forest:

| **Sample type** | **No. of samples** | **No. *Ab* positive samples** | **Percentage positive samples** |
|---|---|---|---|
| **Compost, Wernigerode (Germany)** | **174** | **98** | **56.3** |
| **Forest, Wernigerode (Germany)** | **269** | **2** | **0.7** |

#### Cat-captured mammals (Germany)

| **Species** | **No. samples** | **No. *Ab* positive samples** | **Percentage positive samples** |
|---|---|---|---|
| Rodents | 154 | 3 | 2.0 |
| Shrew | 64 | 3 | 4.7 |
| Other mammals (mole, weasel) | 2 | 0 | 0 |

#### Rats & other Rodents (wildlife, captured and laboratory, Germany)

| **Species** | **No. samples** | **No. *Ab* positive samples** | **Percentage positive samples** |
|---|---|---|---|
| Rats (wildlife, captured, laboratory) | 491 | 0 | 0 |
| Diverse Rodents (*Microtus*, *Myodes*, *Apodemus* and *Arvicola*, wildlife) | 532 | 0 | 0 |

#### Miscellaneous

Diverse pellets and feathers of undefined origin:

6 Pellets, putatively from *Tyto alba* and *Asio otus*, respectively: 2 isolates (33%)

#### Overview: Distinct* *A. baumannii* isolated from outside hospitals (2013-2021)

| | |
|---|---|
| Stork-associated | 926 (837 from nestlings, 53 from pellets, 36 div.) |
| Other animals (diverse, excl. earthworms) | 86 |
| Earthworms | 40 |
| Soil (including compost) | 278 |
| Plant rhizosphere | 78 |
| Environmental (diverse including water & air) | 44 |
| **Sum** | **1,452** |

* From individual samples, only isolates with different OXA-51-like variants were considered distinct. Since individual samples yielded up to 4 distinct isolates, the number of positive samples is not identical to the number of isolates.

**<u>Supplementary Table S2:</u> Blood parameters of white stork nestlings colonized or not with *A. baumannii***

| Blood parameters | Mean value | SD | t | p |
|---|---|---|---|---|
| Packed cell volume (PCV) + | 30,11 | 2,43 | 0,506 | 0,616 |
| Packed cell volume (PCV) - | 29,93 | 2,29 | | |
| Hemoglobin concentration (HGB) + | 8,43 | 0,93 | -8,138 | **0** |
| Hemoglobin concentration (HGB) - | 8,36 | 1,22 | | |
| Red blood cell count (RBC) + | 1,36 | 0,17 | -2,106 | **0,043** |
| Red blood cell count (RBC) - | 1,43 | 0,16 | | |
| White blood cell count (WBC) + | 19,87 | 0,02 | 73,975 | **0** |
| White blood cell count (WBC) - | 18,55 | 0,02 | | |
| Mean corpuscular hemoglobin (MCH) + | 62,7 | 9,93 | 3,163 | **0,003** |
| Mean corpuscular hemoglobin (MCH) - | 58,65 | 7,87 | | |
| Mean corpuscular volume (MCV) + | 4,71 | 0,22 | 2,865 | **0,007** |
| Mean corpuscular volume (MCV) - | 4,59 | 0,22 | | |

"+" and "-" indicate groups of white stork nestlings colonized or not with *A. baumannii*

**Blood sample collection and processing**

Two ml of blood was collected from each individual from the basilic vein. Due to the delicate walls of blood vessels in birds, it was decided to use a combination kit consisting of a 20G intravenous cannula and a 10 ml syringe. The blood from the syringe was then poured into the K2-EDTA tubes to prevent the blood from clotting until laboratory analysis was conducted. Blood sample analysis included: red blood cell number (RBC), haematocrit (PCV), and hemoglobin content (HGB). To determine the haematocrit, the blood was transferred to the capillaries, without anticoagulant, with a volume of 75 µl. The capillaries were centrifuged for 3 minutes at 12,000 RPM. The result was read on a hematocrit reader (MPW) (Szutowicz and Raszeja-Specht 2009; Turgeon 2004). Drabkin's cyanomethemoglobin method (Drabkin 1945) was used to determine hemoglobin in nestlings. For this purpose, 20 µl of blood was added to a test tube containing 5 ml of Drabkin's reagent (Stamar), thoroughly shaken and allowed to stand for 20 minutes. After this time, the tubes were centrifuged at 3000 RPM for 5 minutes to separate the membranes and cell nuclei of the hemolyzed erythrocytes. Supernatant (hemoglobin bound to Drabkin's reagent) was poured into standard cuvettes (MedLab products) of a plastic spectrophotometer. Then, using a UV-VIS spectrophotometer ($\lambda$ = 540 nm) (Prove 300, Merck), the absorbance of the solution against cyanomethemoglobin was measured.

The red blood cell count was counted using a Bürker hemocytometer, with Natt and Herrick staining being used because of the presence of nuclei in the erythrocytes (Natt and Herrick 1952). For this purpose, 20 µl of blood (dilution 1:200) was added to a test tube containing 4 ml of dye. The tubes were stoppered and continuously mixed with a roller mixer for 15 minutes. After this time, 20 µl of the mixture was placed in the hemocytometer chamber and allowed to stabilize for 2 minutes. Red blood cells were counted in 80 small squares at 400x magnification (MT5300, Meiji).

The number of white blood cells was determined analogously to the number of red blood cells. The only difference was that blood cells were counted from the entire area of the Bürker counting chamber. The same magnification was also used - 400x (MT5300, Meiji).

Mean corpuscular hemoglobin (MCH) and mean corpuscular volume (MCV) was calculated based on red blood cell number, haematocrit, and hemoglobin content values (Campbell 1995).

#### Discussion

With an insignificant difference in packed cell volume, a significantly lower RBC value would indicate that individuals colonized with *A. baumannii* were losing water from the blood plasma. Differences in the number of red blood cells, despite statistically significant, are not very high, so it can be that individuals colonized with *A. baumannii* were more likely to become dehydrated. Research in this direction should be conducted in the future.
An increase in MCV (mean corpuscular volume) in individuals colonized with *A. baumannii* indicates an increased absorption of blood plasma water by red blood cells. At the same time, colonized individuals were characterized by a higher concentration of hemoglobin in the blood and, consequently, also a higher content of hemoglobin in the blood cell (MCH). The increase in the concentration of HGB and MCH enables an increase in the intensity of metabolic changes that enable an increased reaction of the immune system. Those values of red blood parameters were different from other bird species with infections (Davis et al. 2004, Lanzarot et al. 2005, Santos et. al. 2006).
No increased mortality of individuals infected with the bacterium was observed. Despite the young age of the studied storks, it can be suggested that they already have developed defense mechanisms. A statistically significant increase in the number of white blood cells (WBC) in individuals colonized with *A. baumannii* could be a reaction of the immune system to this bacterium indicating infection rather than mere colonization.
On the part of the red and white blood cell system, juvenile white storks appear prepared for the threats associated with infection by bacteria such as *A. baumannii*.

**Suppl. Table S6: Remarkable relationship between strains described in this study and strains of public databases as well as among strains described in this study**

| **Our collection** | **NCBI (or other source as stated)** | **SNP distance (core genome set)** | **Pairwise comparison** |
|---|---|---|---|
| 14-2PW1 (Poland, 2014, white stork, OXA-433) | GCA_900494865 (Thailand, 2016, human, OXA-433) | 35 | Shared genes: 3338 SNPs: 143 |
| 17-Zw_133-2 (Germany, 2017, earthworm, OXA-104) | GCA_008985405 (Japan, 2012, human, OXA-104) | 609 | Shared genes: 3442 SNPs: 4005 |
| 17-HoPe_S7-1 (Germany, 2017, soil, OXA-51) | ERR1226902 (Germany, 2005, human, OXA-51, IC4) | 3958 | n.d. |
| 14-73D4 (Poland, 2014, white stork, OXA-402) | GCA_016473355 (USA, 2019, human, OXA-402) | 2714 | Shared genes: 3159 SNPs: 9375 |
| 19-Pos11 (Poland, 2019, white stork, OXA-180) | GCA_001261895 (France, 2013, human, OXA-none) | 180 | Shared genes: 3299 SNPs: 1598 |
| 18-48P05L-2 (Poland, white stork, OXA-762) | GCA_008988845 (Japan, 2013, human, OXA-762) | 48 | Shared genes: 3325 SNPs: 1057 |
| 15-PosEisch11-1 (Poland, 2015, white stork, OXA-132) | GCA_003057525 (Lebanon, 2017, human, OXA-132) | 489 | Shared genes: 3305 SNPs: 3043 |
| 17-13P88K-1 (Poland, 2017, white stork, OXA-762) | K19M21 (Germany, 2019, human, OXA-762) | 1286 | Shared genes: 3577 SNPs: 3895 |
| 13-156-2C (Poland, 2013, white stork, OXA-379) | 18-Pos692-1 (Poland, 2018, white stork, OXA-379) | 4 | Shared genes: 3499 SNPs: 244 |
| 15-7P651-1 (Poland, 2015, white stork, OXA-64) | GCA_004354185 (Canada, 2013, human, OXA-64) | 27 | Shared genes: 3439 SNPs: 951 |
| K19M5 (Germany, 2019, human, OXA-781) | K19-K5 (Germany, human, OXA-781) | 2 | n.d. |
| 17-ZW_S24.2-2 (Germany, 2017, soil, OXA-716) | GCA_000581995 (USA, human, OXA-716) | 1510 | Shared genes: 3323 SNPs: 5350 |
| 15-LoGeIst3-1 (Germany, 2015, white stork, OXA-378) | GCA_008984505 (Japan, 2013, human, OXA-378) | 1367 | n.d. |
| 30813_S1 (Germany, 2013, human, OXA-378) | GCA_002573905 (Germany, 2016, goose, OXA-378) | 78 | Shared genes: 3385 SNPs: 1357 |
| 18-Pos121-2 (Poland, 2018, white stork, OXA-714) | K19M2 (Germany, 2019, human, OXA-714) | 869 | Shared genes: 3357 SNPs: 4621 |
| 18-CSP68-2 (Poland, 2018, plant root, OXA-761) | GCA_001721425 (ATCC 17978) | 4403 | Shared genes: 3245 SNPs: 9642 |
| 18-31P37O-2 (Poland, 2018, white stork, OXA-262) | GCA_000369265 (CZ, 1992, human, OXA-262, NIPH 67) | 427 | n.d. |
| 16-Klo_50-1 (Poland, 2016, white stork, OXA-64) | GCA_009001915 (Japan, 2006, human, OXA-64, IC7) | 2806 | n.d. |

Supplementary Table S6

| | | | |
|---|---|---|---|
| U17-HoPeP5-1 (Germany, 2017, plant root, OXA-769) | 32503-Wdh_S3 (Germany, 2013, dog, OXA-769) | 4 | Shared genes: 3496<br>Accessory genes: 43!<br>SNPs: 17 |
| 19-Spa223 (Spain, 2019, white stork, OXA-121) | GCA_003070915 (China, OXA-121) | 1230 | n.d. |
| 15-Klo_47-1 (Poland, 2015, white stork, OXA-65) | GCA_016609745 (USA, 2003, human, OXA-65, IC5) | 1465 | n.d. |
| 17-HoPe_P2-2 (Germany, 2017, plant root, OXA-106 | GCA_000580815 (USA, human, OXA-106) | 102 | Shared Genes: 3356<br>SNPs: 1769 |
| 17-ZW108-1 (Germany, 2017, earthworm, OXA-770) | GCA_004354145 (Canada, 2013, human, OXA-770) | 120 | n.d. |
| 15-220-51-2 (Poland, 2015, white stork, OXA-689) | GCA_016505755 (Italy, 2018, human, OXA-689) | 213 | n.d. |
| 16-Klo37-1 (Poland, 2016, white stork, OXA-106) | GCA_000505685 (China, OXA-106) | 2167 | n.d. |
| 16-Klo_52-3 (Poland, 2016, white stork, OXA-51) | GCA_016485765 (South Africa, 2014, OXA-none) | 2059 | n.d. |
| 16-Klo_26-1 (Poland, 2016, white stork, OXA-69) | GCA_001674695 (2014, human, OXA-69) | 1113 | n.d. |
| 15-22064-2 (Poland, 2015, white stork, OXA-374) | 19-Spa420-1 (Spain, 2019, stork, OXA-374) | 108 | Shared genes: 3379<br>SNPs: 2540 |
| 18-Pos111-1 (Poland, 2018, white stork, OXA-374) | GCA_006493955 (Germany, 2009, human, OXA-374) | 5332 | n.d. |
| 17-HoPe_P1-2 (Germany, 2017, plant root, OXA-68) | GCA_016511655 (Australia, 2011, human, OXA-68, IC8);<br>IMT30938 (this study, Germany, 2013, human, OXA-68) | 3803<br><br>85 | n.d.<br><br>n.d. |
| 15-7P830-2 (Poland, 2015, white stork, OXA-853) | GCA_900495975 (Thailand, 2016, human, OXA-853) | 681 | n.d. |
| 17-33P33T-1 (Poland, white stork, OXA-739) | GCA_900495605 (Thailand, 2016, human, OXA-739) | 1511 | n.d. |
| 15-LoGeW2-11-2 (Germany, 2015, stork) | 15-Posen_P44-1 (Poland, 2015, stork) | 256 | n.d. |
| 17-HoPe_S6-1 (Germany, 2017, soil, OXA-51) | 16-Klo_103-1 (Poland, 2016, white stork, OXA-51) | 5 | Shared genes: 3402<br>Accessory genes: 106<br>SNPs: 21 |
| 17-ZW-P5-2 (Germany, 2017, plant root, OXA-343) | U17-HoPe-P3-3 (Germany, 2017, plant root, OXA-343) | 1 | Shared Genes: 3498<br>Accessory genes: 151<br>SNPs: 61 |
| 15-22062-1 (Poland, 2015, white stork, OXA-343) | GCA_002367895 (Luxembourg, 2016, grey parrot, OXA-343) | 736 | n.d. |
| T20-Cu8 (Germany, 2020, human, OXA-390) | 13-658C (Poland, 2013, white stork, OXA-390) | 1414 | n.d. |
| 17-Lo_4-1 (Germany, 2017, white stork, OXA-51) | GCA_001693195 (Poland, white stork, OXA-51, strain 29D2) | 8 | Shared genes: 3472<br>Accessory genes: 143<br>SNPs: 1497 |

**Supplementary Table S8**

**Further evidence for the interrelatedness of the *A. baumannii* life cycle with the fungal world**

- *A. baumannii* isolate from strawberry rhizosphere has antagonistic activity against *Verticillium* fungi (1).
- *A. baumannii* use DNase secreted via a type VI secretion system to kill fungi (2)
- Chemical warfare between *Aspergillus* and *Acinetobacter*: Aspergillomarasmine overcomes metallo-β-lactamase resistance of *Acinetobacter* and other Gram-negatives (3)
- *Acinetobacter* contribute to decomposition of insect cadavers after fungal infection (4)
- Endosymbiotic *Acinetobacter* were discovered in the entomopathogenic fungus *Pandora neoaphidis* (5)
- During maturation of Camembert cheese, it is only after growth of the yeast *Debaryomyces hansenii* that colonization with *Acinetobacter* spp. follows (6)
- Antifungal volatile compounds produced by *A. baumannii* are highly effective to suppress growth of *Aspergillus flavus* (7)
- Co-occurrence of *Acinetobacter* and *Aspergillus* during production of sufu, a fermented soybean food (8) and during natural fermentation of milk (9)
- *Acinetobacter* spp. and various fungi are dominant microorganisms found attached to airborne particulates in vegetable plastic greenhouse (10)
- *Penicillium brevicompactum*/*Penicillium expansum* and *Acinetobacter calcoaceticus* isolated from drinking water form mixed biofilms (11)
- *Acinetobacter* isolates including representatives of *A. pittii* from frog skin with antifungal activity (12)
- *A. baumannii* is capable of growing on xylose, arabinose and ribose (13), pentose sugars released during the first steps of decomposition of plant material by extracellular enzymes produced by fungi (14, 15)

#### - References


1. **Berg G, Roskot N, Steidle A, Eberl L, Zock A, Smalla K.** 2002. Plant-dependent genotypic and phenotypic diversity of antagonistic rhizobacteria isolated from different Verticillium host plants. Appl Environ Microbiol **68:**3328-3338.
2. **Luo J, Chu X, Jie J, Sun Y, Guan Q, Li D, Luo ZQ, Song L.** 2023. Acinetobacter baumannii Kills Fungi via a Type VI DNase Effector. mBio doi:10.1128/mbio.03420-22:e0342022.
3. **King AM, Reid-Yu SA, Wang W, King DT, De Pascale G, Strynadka NC, Walsh TR, Coombes BK, Wright GD.** 2014. Aspergillomarasmine A overcomes metallo-β-lactamase antibiotic resistance. Nature **510:**503-506.
4. **Kryukov VY, Kabilov MR, Smirnova N, Tomilova OG, Tyurin MV, Akhanaev YB, Polenogova OV, Danilov VP, Zhangissina SK, Alikina T, Yaroslavtseva ON, Glupov VV.** 2019. Bacterial decomposition of insects post-Metarhizium infection: Possible influence on plant growth. Fungal Biol **123:**927-935.
5. **Chen C, Chen X, Xie T, Louis Hatting J, Yu X, Ye S, Wang Z, Shentu X.** 2016. Diverse bacterial symbionts of insect-pathogentic fungi and possible impact on the maintenance of virulence during infection. Symbiosis **69:**47-58.
6. **Addis E, Fleet GH, Cox JM, Kolak D, Leung T.** 2001. The growth, properties and interactions of yeasts and bacteria associated with the maturation of Camembert and blue-veined cheeses. Int J Food Microbiol **69:**25-36.
7. **Kadhim M.** 2016. In Vitro antifungal potential of Acinetobacter baumannii and determination of its chemical composition by gas chromatography-mass spectrometry. Der Pharma Chemica **8:**657-665.
8. **Xu D, Wang P, Zhang X, Zhang J, Sun Y, Gao L, Wang W.** 2020. High-throughput sequencing approach to characterize dynamic changes of the fungal and bacterial communities during the production of sufu, a traditional Chinese fermented soybean food. Food microbiology **86:**103340.
9. **Xu WL, Li CD, Guo YS, Zhang Y, Ya M, Guo L.** 2021. A snapshot study of the microbial community dynamics in naturally fermented cow's milk. Food Sci Nutr **9:**2053-2065.
10. **Nie C, Geng X, Ouyang H, Wang L, Li Z, Wang M, Sun X, Wu Y, Qin Y, Xu Y, Tang X, Chen J.** 2022. Abundant bacteria and fungi attached to airborne particulates in vegetable plastic greenhouses. Sci Total Environ doi:10.1016/j.scitotenv.2022.159507**:**159507.
11. **Simões LC, Chaves AFA, Simões M, Lima N.** 2023. Interactions between Penicillium brevicompactum/Penicillium expansum and Acinetobacter calcoaceticus isolated from drinking water in biofilm development and control. International Journal of Food Microbiology **384:**109980.
12. **Cevallos MA, Basanta MD, Bello-López E, Escobedo-Muñoz AS, González-Serrano FM, Nemec A, Romero-Contreras YJ, Serrano M, Rebollar EA.** 2022. Genomic characterization of antifungal Acinetobacter bacteria isolated from the skin of the frogs Agalychnis callidryas and Craugastor fitzingeri. FEMS Microbiol Ecol **98**.
13. **Alberti L, König P, Zeidler S, Poehlein A, Daniel R, Averhoff B, Müller V.** 2023. Identification and characterization of a novel pathway for aldopentose degradation in Acinetobacter baumannii. Environ Microbiol doi:10.1111/1462-2920.16471.
14. **Prieto A, de Eugenio L, Méndez-Líter JA, Nieto-Domínguez M, Murgiondo C, Barriuso J, Bejarano-Muñoz L, Martínez MJ.** 2021. Fungal glycosyl hydrolases for

sustainable plant biomass valorization: Talaromyces amestolkiae as a model fungus. Int Microbiol **24:**545-558.

15. **Seiboth B, Metz B.** 2011. Fungal arabinan and L-arabinose metabolism. Appl Microbiol Biotechnol **89:**1665-1673.

**Suppl. Material S1: Supplementary information on GIS-based analyses**

A total of 295 nests were analysed (50 from the southern part of Lubsz voivodeship near Zielona Góra city, hereafter "Zielona Góra"; 95 from the southern part of Greater Poland voivodeship near Leszno city, hereafter "Leszno"; 150 from Opole voivodeship near Opole city, hereafter "Opole"). From these 295 nests during breeding seasons 2015-2018, we analysed 454 broods (mean number of broods per nest was 1.5). A total number of 1080 chicks was analysed.

First, we tested for spatial autocorrelation with Moran's local indicator (Legendre and Legendre 1998). As data set we used geographic position of 295 nests and whether we isolated or not *A. baumannii* in nests during four years of study. We found no spatial autocorrelation (Table S1, Fig. S1). The spatial autocorrelation was computed using SAM software (Rangel et al. 2010).

For each nest we obtained several environmental variables in a radius of 3.5 km and 4.0 km around the nest. We determined the landscape variable according to Corine land cover classification (EEA - European Environment Agency 2006) as proportion to total buffer area: artificial surface (CLC_1), arable land (CLC_2.1), pastures (CLC_2.3), heterogeneous agricultural area (CLC_2.4), forests (CLC_3.1), water reservoirs area (CLC_5.1.2) and water courses length (CLC_5.1.1). The GIS data were proceed by Quantum GIS Software ("Quantum GIS Geographic Information System" 2010). Before analysis, the environmental variables were log-transformed in order to minimize effects of detached observations and skewed distributions.

We checked whether there is multicollinearity between those environmental variables by using variance inflation factor. We found that there is collinearity between arable land and forest areas. There is strong negative correlation between them (Pearson correlation for radius 3.5 km: - 0.72, $p < 0.001$ and for radius 4.0 km: -0.75, $p < 0.001$), so we excluded forest variable from the analysis. So it should be noted when interpreting the results that high area of arable land indicates low area of forest.

Precipitation data were based on shared data from the Institute of Meteorology and Water Management National Research Institute. Yearly sum of rainfall for nests was calculated based on assignment to the closest meteorological station in a 35 km radius from the nest.

We compared the environmental variables from buffers in a radius of 3.5 km around nests. According to this, we compared regions using Kruskal-Wallis tests. The results are shown in Table S2. According to this, the Zielona Góra region exhibited the most extensive agricultural use (mostly pasture). On the other hand, Opole was the region with highest arable area, followed by Leszno and Zielona Góra. The ratio of forests was highest in buffers around nests in Zielona Góra.

Our dependent variable was the proportion of the number of positive chicks to total number of chicks in the broods. We used a generalised linear model with a binomial error distribution. Because the broods measurements were repeated in different years we used a mixed model with nest id as random factor. Also as random variable we used the year of study. As independent variable we used above mentioned environmental variables, precipitation and mean age of fledglings in the broods (age was estimated according to size of beak) and region (as factor variable with three levels: Zielona Góra, Leszno, Opole). We started from the global models with the interaction of region factor with all other environmental covariates. The final model was selected using backward stepwise removal of nonsignificant effects of interactions ($P < 0.05$) (Zuur et al. 2009).

For the statistical analyses we used R software (R Development Core Team 2018) and for mixed modelling approach we used "lme4" package (Bates et al. 2014).

**Results**

The proportion for *A. baumannii* in Zielona Góra was 0.216 ± 0.057 s.e. (stander error), in Leszno region 0.408 ± 0.075 s.e., and in Opole region 0.292 ± 0.061 s.e.. According to the model, we found that regions differ in *A. baumannii* presence. Post-hoc Tukey tests indicate that the Zielona Góra region exhibited a significantly lower proportion than Leszno (OR (odds ratio) = 0.40 ± 0.12 s.e., P < 0.01), but was not significantly different to Opole (OR = 0.667 ± 0.19 s.e., P = 0.311). Comparing the Leszno to the Opole region there was a higher proportion of *A. baumannii*, however it was not significant (OR = 1.668± 0.41, P = 0.094).

We found a significant interaction between region and precipitation variable (Table 1). We found that the Leszno region differs significantly from the Zielona Góra region, while no significant difference was observed between Opole and Zielona Góra regions (Fig. 1A). We found a positive relation of *A. baumannii* presence with precipitation for Leszno region (P < 0.01) but there was a negative relation with this variable in Zielona Góra (P = 0.028) and Opole (P < 0.001).

Our analysis revealed that the mean age of fledglings was significant (Table 1). We found a negative relation between the mean age of fledglings and *A. baumannii* presence (Fig 1B). We found a positive relation with artificial surface and heterogeneous agriculture area in a radius of 3.5 km around nests (Table 1, Fig 1C and 1D). In case of area in a 4 km radius, only artificial surface was significant (Table 1).

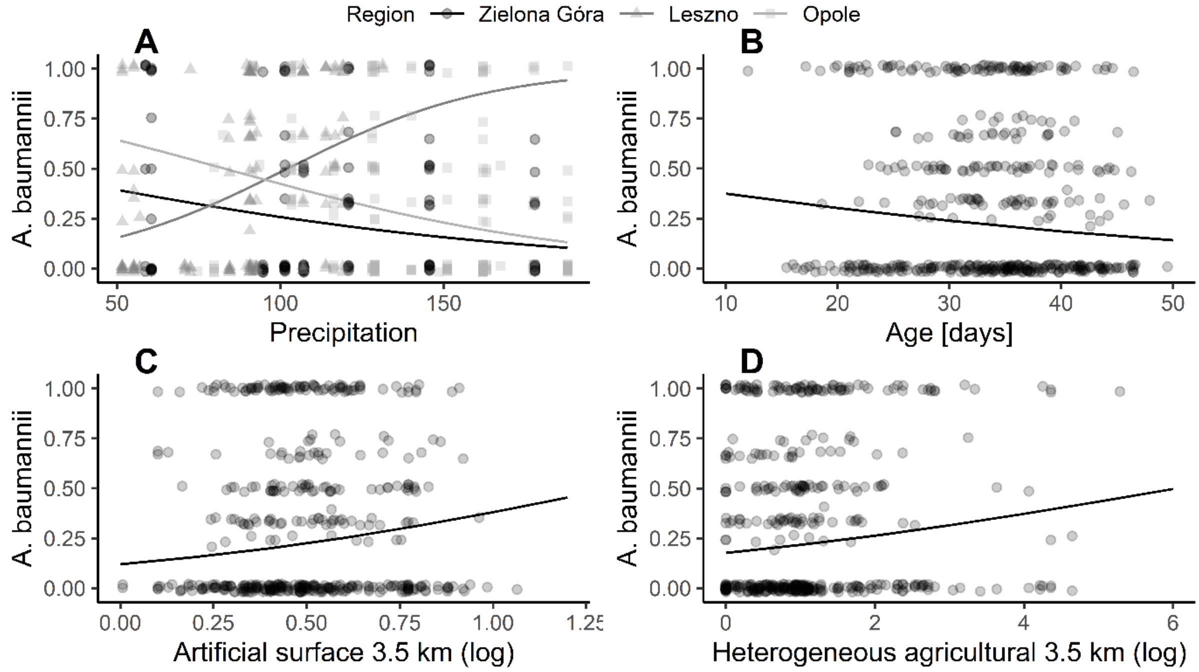


Fig. 1. Prediction of fitted model showing relationship between different explanatory variables and *A. baumannii* presence in the brood of White Stork (*Ciconia ciconia*). Panel A shows significant interaction between region and precipitation. Panels B-D show additive effects of age, artificial surface and heterogeneous agriculture on *A. baumannii* occurrence, respectively.

Table 1. The estimated effects of two models, where of environmental variables was counted differently – from buffers of radius 3.5 km and 4.0 km around the nest. We used GLMM with binomial error distribution and two random effects (r), where dependent variable was the proportion of fledglings positively diagnosed for *A. baumannii* to the total number of fledglings. P < 0.05 are bolded.

| | buffers 3.5 km | | | | buffers 4.0 km | | | |
|---|---|---|---|---|---|---|---|---|
| | Est | SE | Z | P | Est | SE | Z | P |
| (Intercept) | 0.373 | 1.435 | 0.26 | 0.795 | 0.617 | 1.556 | 0.396 | 0.692 |
| Water bodies | -0.665 | 0.859 | -0.775 | 0.439 | -0.747 | 0.733 | -1.02 | 0.308 |
| Artificial surfaces | 1.507 | 0.647 | 2.327 | **0.020** | 1.413 | 0.605 | 2.336 | **0.019** |
| Arable | -0.060 | 0.773 | -0.077 | 0.939 | -0.396 | 0.815 | -0.486 | 0.627 |
| Pastures | -0.325 | 0.468 | -0.695 | 0.487 | -0.297 | 0.455 | -0.654 | 0.513 |
| Heterogenagri | 0.254 | 0.117 | 2.179 | **0.029** | 0.139 | 0.097 | 1.435 | 0.151 |
| Water courses | -0.214 | 0.344 | -0.622 | 0.534 | -0.149 | 0.321 | -0.463 | 0.643 |
| Region:Leszno | -3.452 | 1.031 | -3.349 | **0.001** | -3.228 | 1.026 | -3.146 | **0.002** |
| Region:Opole | 1.236 | 0.937 | 1.318 | 0.187 | 1.490 | 0.937 | 1.59 | 0.112 |
| Precip | -0.013 | 0.006 | -2.063 | **0.039** | -0.012 | 0.006 | -1.92 | 0.055 |
| Age | -0.034 | 0.015 | -2.307 | **0.021** | -0.033 | 0.015 | -2.242 | **0.025** |
| Dist residence | 0.401 | 0.248 | 1.616 | 0.106 | 0.419 | 0.248 | 1.689 | 0.091 |
| Dist non-resid | -0.085 | 0.219 | -0.391 | 0.696 | -0.076 | 0.218 | -0.35 | 0.726 |
| RegionLeszno: precip | 0.045 | 0.010 | 4.487 | **<0.001** | 0.044 | 0.010 | 4.395 | **<0.001** |
| RegionOpole: precip | -0.006 | 0.007 | -0.836 | 0.403 | -0.007 | 0.007 | -0.984 | 0.325 |
| r(Nest_ID) | 1.238 | 1.113 | | | 1.226 | 1.107 | | |
| r(Year) | 0.084 | 0.289 | | | 0.080 | 0.283 | | |

**Abbreviations: Dist residence**, distance to residential building; **Dist non-resid**, distance to non-residential building; **precip**, precipitation.

Table S1. Tests for spatial autocorrelation with Moran's local indicator (I). Minimal value of this indicator is -1 and means perfect dispersion, 0 means random dispersion and high cluttering would be close to 1. Dist Cntr – distance classes (in meters)

| DistCntr | Moran's I | P | I (max) | I/I(max) |
|---|---|---|---|---|
| 11255.1 | 0.026 | 0.111 | 0.341 | 0.075 |
| 33765.31 | 0.001 | 0.94 | 0.216 | 0.005 |
| 56275.52 | 0.008 | 0.508 | 0.375 | 0.022 |
| 78785.73 | 0.004 | 0.834 | 0.557 | 0.007 |
| 101295.9 | -0.024 | 0.196 | 0.61 | -0.04 |
| 123806.2 | -0.019 | 0.362 | 0.338 | -0.055 |
| 146316.4 | -0.042 | 0.045 | 0.37 | -0.114 |
| 168826.6 | -0.005 | 0.688 | 0.229 | -0.024 |
| 191336.8 | -0.017 | 0.126 | 0.213 | -0.079 |
| 213847 | -0.007 | 0.533 | 0.209 | -0.036 |
| 236357.2 | 0.017 | 0.327 | 0.326 | 0.052 |
| 258867.4 | -0.004 | 0.889 | 0.514 | -0.007 |
| 281377.6 | -0.022 | 0.266 | 0.326 | -0.066 |
| 303887.8 | 0.002 | 0.889 | 0.493 | 0.004 |
| 326398 | <.001 | 0.97 | 0.658 | <.001 |

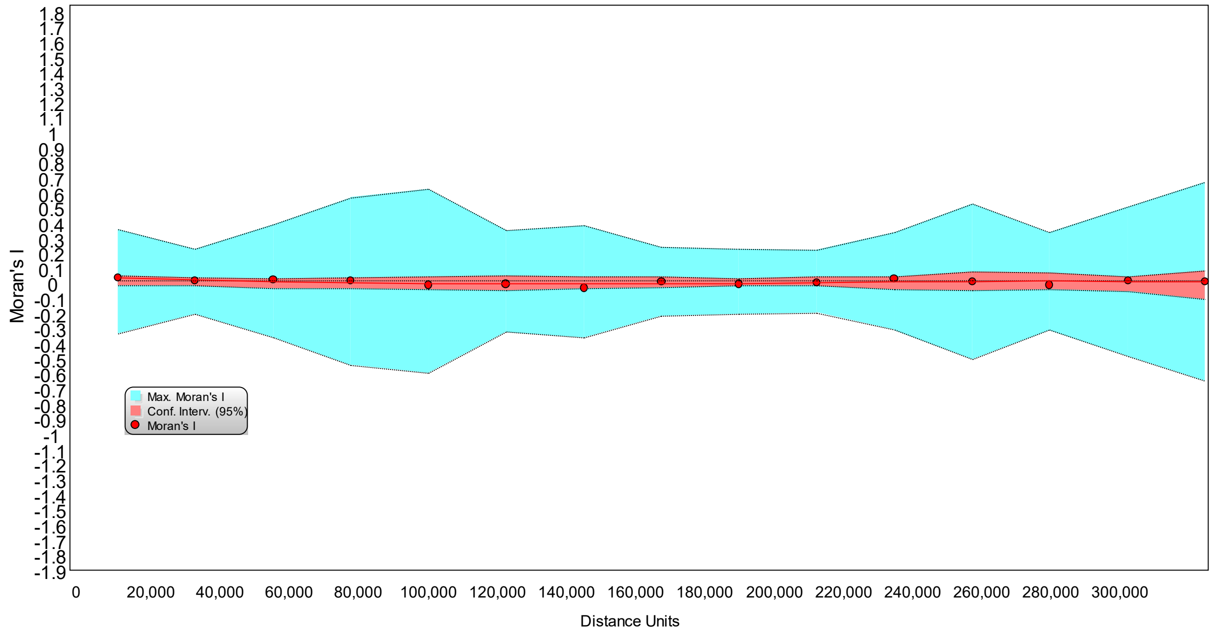


Fig. S1. Spatial correlogram

Table S2. Kruskal-Wallis tests and multiple comparisons between regions.

| | Min. | 1st Qu. | Median | Mean | 3rd Qu. | Max. | | |
|---|---|---|---|---|---|---|---|---|
| **Artificial surfaces** | | | | | | | H = 5.299, p = 0.07071 | |
| Leszno | 0.7 | 3.1 | 4.9 | 5.8 | 7.0 | 22.5 | Leszno vs Zielona Góra | 0.065 |
| Opole | 0.9 | 3.7 | 5.5 | 6.2 | 7.4 | 27.5 | Opole vs Leszno | 0.230 |
| Zielona Góra | 0.0 | 4.0 | 7.3 | 7.9 | 12.8 | 17.1 | Zielona Góra vs Opole | 0.271 |
| **Arable** | | | | | | | H = 63.281, p < 0.001 | |
| Leszno | 17.9 | 49.8 | 59.9 | 58.3 | 70.0 | 85.8 | Leszno vs Zielona Góra | <0.001 |
| Opole | 16.3 | 52.8 | 69.3 | 65.4 | 82.0 | 93.1 | Opole vs Leszno | <0.001 |
| Zielona Góra | 10.5 | 20.2 | 42.4 | 38.3 | 43.6 | 81.2 | Zielona Góra vs Opole | <0.001 |
| **Pastures** | | | | | | | H = 77.888 , p < 0.001 | |
| Leszno | 0.7 | 9.0 | 14.7 | 17.9 | 21.9 | 69.3 | Leszno vs Zielona Góra | <0.001 |
| Opole | 0.0 | 3.0 | 6.2 | 6.9 | 10.2 | 30.4 | Opole vs Leszno | <0.001 |
| Zielona Góra | 0.0 | 9.8 | 17.5 | 14.3 | 18.5 | 29.7 | Zielona Góra vs Opole | <0.001 |
| **Heterogeneous agriculture areas** | | | | | | | H = 23.738, p < 0.001 | |
| Leszno | 0.0 | 0.5 | 1.7 | 1.9 | 3.1 | 6.0 | Leszno vs Zielona Góra | <0.001 |
| Opole | 0.0 | 1.5 | 2.8 | 3.5 | 4.8 | 13.7 | Opole vs Leszno | <0.001 |
| Zielona Góra | 0.0 | 2.4 | 2.6 | 3.2 | 4.2 | 11.3 | Zielona Góra vs Opole | <0.001 |
| **Forests** | | | | | | | H = 32.511, p < 0.001 | |
| Leszno | 0.1 | 7.7 | 13.2 | 15.2 | 20.7 | 45.0 | Leszno vs Zielona Góra | <0.001 |
| Opole | 0.0 | 3.8 | 11.8 | 17.4 | 26.7 | 73.6 | Opole vs Leszno | <0.001 |
| Zielona Góra | 5.7 | 17.5 | 23.5 | 33.5 | 53.1 | 78.6 | Zielona Góra vs Opole | <0.001 |
| **Water bodies** | | | | | | | H = 15.781, p <0.001 | |
| Leszno | 0.0 | 0.0 | 0.0 | 0.9 | 1.0 | 8.0 | Leszno vs Zielona Góra | 0.699 |
| Opole | 0.0 | 0.0 | 0.0 | 0.5 | 0.0 | 13.4 | Opole vs Leszno | <0.001 |
| Zielona Góra | 0.0 | 0.0 | 0.0 | 0.2 | 0.3 | 3.1 | Zielona Góra vs Opole | 0.022 |
| **Water courses** | | | | | | | H = 143.83, p < 0.001 | |
| Leszno | 0.0 | 0.0 | 0.0 | 0.0 | 0.0 | 0.0 | Leszno vs Zielona Góra | <0.001 |
| Opole | 0.0 | 0.0 | 0.0 | 0.1 | 0.0 | 1.3 | Opole vs Leszno | <0.001 |
| Zielona Góra | 0.0 | 0.1 | 1.2 | 0.9 | 1.4 | 1.4 | Zielona Góra vs Opole | <0.001 |
| **Distance to residential buildings** | | | | | | | H = 2.768, p = 0.251 | |
| Leszno | 0.0 | 1.0 | 1.3 | 1.2 | 1.5 | 2.6 | Leszno vs Zielona Góra | 0.703 |
| Opole | 0.0 | 0.9 | 1.2 | 1.2 | 1.5 | 2.8 | Opole vs Leszno | 0.426 |
| Zielona Góra | 0.7 | 1.2 | 1.3 | 1.3 | 1.5 | 1.9 | Zielona Góra vs Opole | 0.305 |
| **Distance to non-residential buildings** | | | | | | | H = 5.0174, p = 0.813 | |
| Leszno | 0.0 | 0.8 | 1.2 | 1.0 | 1.3 | 2.2 | Leszno vs Zielona Góra | 0.078 |
| Opole | 0.0 | 0.9 | 1.2 | 1.1 | 1.4 | 2.8 | Opole vs Leszno | 0.215 |
| Zielona Góra | 0.0 | 1.0 | 1.3 | 1.2 | 1.5 | 2.0 | Zielona Góra vs Opole | 0.331 |
